# Transcriptome dynamics at the *Arabidopsis* graft junction reveal an inter-tissue recognition mechanism that activates vascular regeneration

**DOI:** 10.1101/198598

**Authors:** Charles W Melnyk, Alexander Gabel, Thomas J Hardcastle, Sarah Robinson, Shunsuke Miyashima, Ivo Grosse, Elliot M Meyerowitz

## Abstract

The ability for cut tissues to join together and form a chimeric organism is a remarkable property of many plants, however, grafting is poorly characterized at the molecular level. To better understand this process we monitored genome-wide temporal and spatial gene expression changes in grafted *Arabidopsis thaliana* hypocotyls. Tissues above and below the graft rapidly developed an asymmetry such that many genes were more highly expressed on one side than the other. This asymmetry correlated with sugar responsive genes and we observed an accumulation of starch above the graft that decreased along with asymmetry once the sugar-transporting vascular tissues reconnected. Despite the initial starvation response below the graft, many genes associated with vascular formation were rapidly activated in grafted tissues but not in cut and separated tissues indicating that a recognition mechanism activated that was independent of functional vascular connections. Auxin which is transported cell-to-cell, had a rapidly elevated response that was symmetric, suggesting that auxin was perceived by the root within hours of tissue attachment to activate the vascular regeneration process. A subset of genes were expressed only in grafted tissues, indicating that wound healing proceeded via different mechanisms depending on the presence or absence of adjoining tissues. Such a recognition process could have broader relevance for tissue regeneration, inter-tissue communication and tissue fusion events.

## INTRODUCTION

For millennia people have cut and rejoined plants through grafting. Generating such chimeric organisms combines desirable characteristics from two plants, such as disease resistance, dwarfing and high yields, or can propagate plants and avoid the delays entailed by a juvenile state (1). Agriculturally, grafting is becoming more relevant as a greater number of plants and species are grafted to increase productivity and yield (2). However, our mechanistic understanding of grafting and the biological processes involved, including wound healing, tissue fusion and vascular formation, remain limited.

Plants have efficient mechanisms to heal wounds and cuts, in part through the production of wound-induced pluripotent cells termed callus. The callus fills the gap or seals the wound, and later, differentiates to form epidermal, mesophyll and vascular tissues (3). In grafted *Arabidopsis* hypocotyls, tissues adhere 1-2 days after grafting and the phloem, the tissue that transports sugars and nutrients, connects after three days (4, 5). The xylem, tissue that transports water and minerals, connects after seven days (4). Plant hormones are important regulators of vascular formation, and at the graft junction, both auxin and cytokinin responses increase in the vascular tissue (4–6). Auxin is important for differentiation of vascular tissues whereas cytokinin promotes vascular stem cells, termed cambium, to divide and proliferate in a process known as secondary growth (7, 8). Auxin is produced in the upper parts of a plant and moves towards the roots via cell-to-cell movement. Auxin exporters, including the PIN proteins, transport auxin into the apoplast, whereas auxin importers, such as the AUX and LAX proteins, assist with auxin uptake into adjacent cells (8). Disrupting this transport, such as by mutating *PIN1*, inhibits healing of a wounded stem (9). Blocking auxin transport with the auxin transport inhibitor TIBA (2,3,5-triiodobenzoic acid) in the shoot inhibits vascular formation and cell proliferation at the *Arabidopsis* graft junction (6). In addition to auxin, other compounds, including sugars, contribute to vascular formation. The localised addition of auxin to callus induces phloem and xylem but requires the presence of sugar (10, 11). In plants, sugars are produced in the leaves and transported through the phloem to the roots (12). The role of sugars in vascular formation and wound healing is not well established, however, sugars promote cell division and cell expansion (13), processes important for development including vascular formation.

The molecular and cellular mechanisms for wound healing, tissue reunion and graft formation remain largely unknown. One emerging theme is that the top and bottom of the cut do not behave similarly. Such tissue asymmetry occurs in other plant tissues, most notably leaves. Developing leaf primordia have an inherent asymmetry that is thought to derive from positional signals from the shoot apical meristem that specifies differences between the top and the bottom of the leaf. The molecular mechanisms that establish asymmetry are not well described, though one hypothesis is that auxin contributes towards or acts on the putative signal (14). Asymmetry also appears in cut *Arabidopsis* inflorescence stems where the transcription factor *RAP2.6L* expresses exclusively below the cut whereas the transcription factor *ANAC071* expresses exclusively above the cut (9). Both were important for stem healing and *ANAC071* and a close homologue, *ANAC096*, were important for graft formation (6). Asymmetry also exists in genetic requirements, since *ALF4* and *AXR1*, two genes involved in auxin perception, are important below but not above the graft junction for phloem connection (4). However, *ANAC071* is expressed symmetrically around the hypocotyl graft junction three days after grafting (6) so the extent of asymmetry and the mechanistic basis for it during wound healing remains largely uncharacterised.

Previous efforts have characterised wound healing and tissue reunion using transcriptomic analyses. Mechanical wounding altered approximately 8% of the *Arabidopsis* transcriptome and showed a high degree of overlap with transcriptomic changes elicited by pathogen attack and abiotic stress (15). Stem wounding and wound-induced callus formation altered the expression of hundreds or thousands of genes (9, 16, 17), whereas grafting grape vines, lychee trees and hickory trees induced hundreds or thousands of differentially expressed genes involved in hormone response, wound response, metabolism, cell wall synthesis and signal transduction (5, 18–21). These grafting studies provide limited information, as tissues from above and below the graft junction were not isolated to test whether these tissues behaved differently, and controls were not performed to distinguish how grafting and tissue fusion might differ from a response associated with cut tissues that remained separated. Here, we perform an in-depth analysis to describe the spatial and temporal transcriptional dynamics that occur during healing of cut *Arabidopsis* tissues that are joined (grafted) or left unjoined (separated). We find that the majority of genes differentially expressed are initially asymmetrically expressed at the graft junction and that many of these genes are sugar responsive, which correlates with severing of the phloem tissue and the accumulation of starch above the junction. However, genes associated with cell division and vascular formation activate on both sides of the graft and, similarly, auxin responsiveness activates equally on both sides. We propose that the continuous transport of substances, including auxin, independent of functional vascular connections, promoted division and differentiation, while the enhanced auxin response and blocked of sugar transport provided a unique physiological condition to activate genes specific to graft formation that promote wound healing.

## RESULTS

### Genes are asymmetrically expressed around the graft

Previous analyses revealed an asymmetry in several genes important for tissue reunion or graft formation and identified a number of genes expressed above a cut that were not expressed below (4, 9). To investigate whether asymmetry was a common feature of grafting and tissue reunion, we generated RNA deep sequencing libraries from *Arabidopsis thaliana* hypocotyl tissues immediately above and immediately below the graft junction 0, 6, 12, 24, 48, 72, 120, 168 and 240 hours after grafting (HAG)(Figure 1A). Prior to RNA extraction, we separated top and bottom tissues at the graft junction. We found that the strength required to break apart the graft junction increased linearly (Figure S1) similarly to previously reported breaking strength dynamics of grafted *Solanum pennellii* and *Solanum lycopersicum* (22, 23). When pulling apart grafts to separate top and bottom for sample preparation, grafts broke cleanly with minimal tissue from one half present in the other half (Figure S1, Movie S1, S2). We measured the amount of tissue from tops adherent to bottoms and *vice versa* (Figure S1) and found less than 4% cross-contamination. In addition to grafting, we also prepared libraries from ungrafted hypocotyls (“intact” treatment) and cut plants that had not been reattached (“separated” treatment)(Figure 1A). We herein refer to tissues harvested above the graft junction or from the shoot side of separated tissue as “top” and that from below the graft or from the root side of separated tissue as “bottom” (Figure 1A).

**Figure 1.**
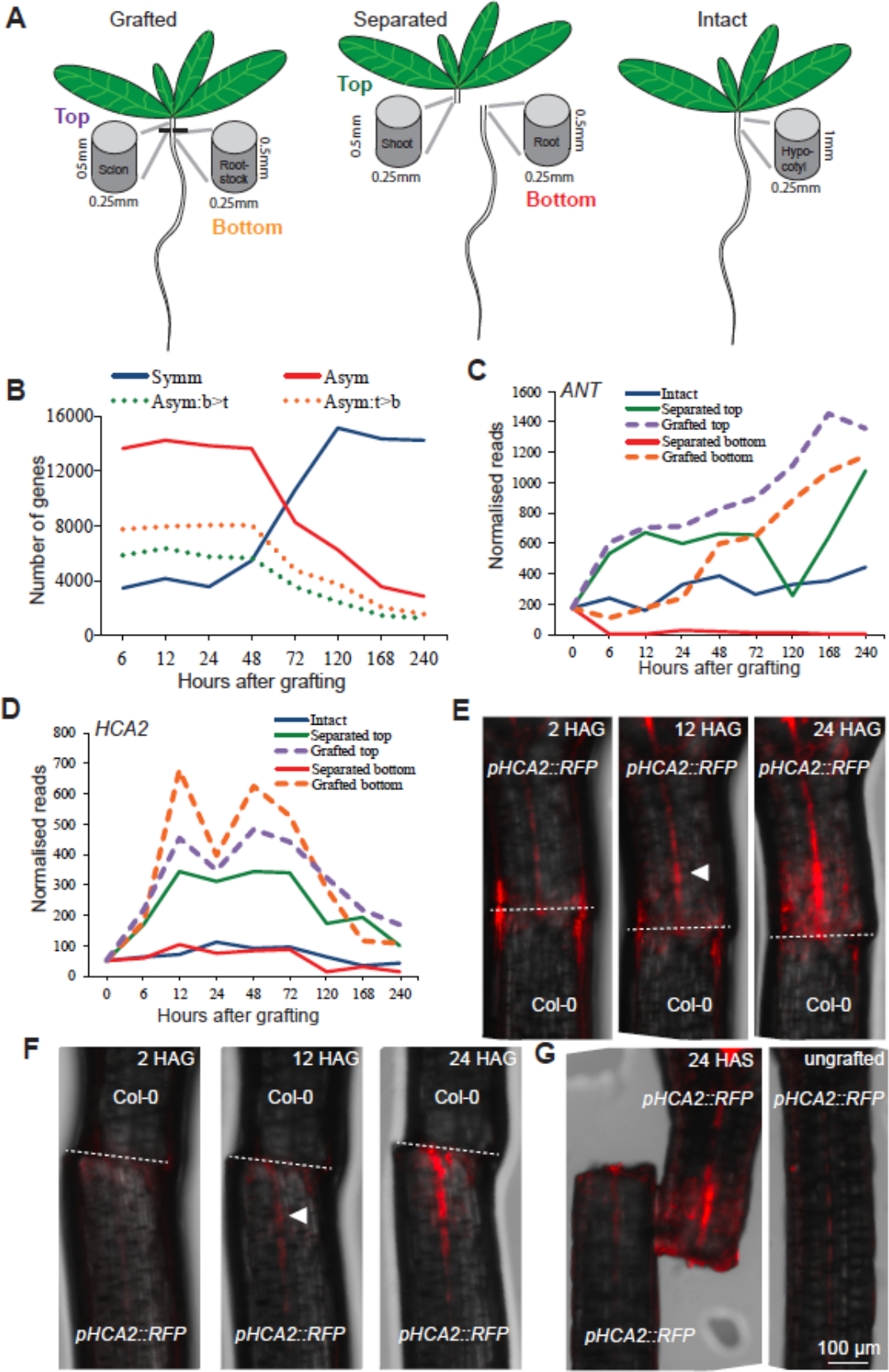
Grafting rapidly activates genes in both asymmetric and symmetric patterns. (A) Intact, cut and separated, or cut and grafted *Arabidopsis* tissues were harvested approximately 0.5mm above (top), 0.5mm below (bottom) the cut site or for intact plants 1mm segments spanning the region where cuts were made in grafted and separated plants. (B) Pairwise analysis between the grafted top and grafted bottom identified sets of protein-coding genes symmetrically or asymmetrically expressed at the graft junction, with an FDR < 0.05 and a likelihood of symmetric/asymmetric expression of greater than 50%. Asymmetrically expressed genes were further divided into those whose RNAs were higher in the top (orange dotted line) or higher in the bottom (purple dotted line), again with an FDR < 0.05 and a likelihood of asymmetric expression greater than 50%. (C,D) Expression profiles for transcripts of cambium-associated genes that initially activate symmetrically (*HCA*) or asymmetrically (*ANT*) plotted for intact, separated and grafted samples. (E-G) *HCA2* transcription upregulates above the graft junction (E) and below the graft junction. (F) *pHCA2::RFP* was grafted to Col-0 roots (E) or Col-0 shoots (F) to avoid ambiguity of signal origin at the junction. *HCA2* was also upregulated in separated shoots but not in intact samples or in high levels in separated roots (G). HAG, hours after grafting. HAS, hours after separation. White triangle denotes initial fluorescent signal, dashed lines denote the graft junction.

RNAs that were differentially expressed as a consequence of grafting (compared to intact hypocotyls) equally in tops and bottoms of grafts (symmetrically expressed) or were more highly expressed in one tissue than the other (asymmetrically expressed), were identified by performing a pairwise analysis of the protein-coding transcriptome datasets that were differentially expressed relative to the intact group. Several thousand RNAs were identified that fit either pattern of expression including the transcript of the cambial markers *HCA2* that was induced symmetrically, and *ANT* that was induced asymmetrically (Figure 1B-G, Figure S2). 6 to 48 hours after grafting, the number of graft-differentially expressed genes that were asymmetrically expressed was roughly 3-fold greater than those symmetrically expressed indicating that tissues above the cut changed their expression dynamics relative to below the cut. However, at 72 hours the numbers were nearly equal, and by 120 hours, the number of symmetrically differentially expressed genes was 3-fold greater than those asymmetrically expressed (Figure 1B). As a second approach, we performed a hierarchical clustering analysis that indicated that the grafted top and grafted bottom became most similar after 72 hours (Figure S3), consistent with the symmetry analysis (Figure 1B). Thus, graft healing and tissue reunion promoted a shift from asymmetry to symmetry (Figure 1B).

### Sugar response correlates with asymmetric gene expression

The shift from asymmetry to symmetry could be due to phloem reconnection at 72 hours ((4), Figure 2A) and the resumption of hormone, protein and sugar transport. We tested a role for sugar by grafting in the presence of exogenous sucrose which has previously been reported to affect grafting success (24). Low levels of exogenous sucrose lowered grafting efficiency (Figure 2A), suggesting that differential sugar responses at the graft junction might be important for vascular reconnection. Expression of *ApL3*, a gene whose expression is induced by sugar (25), was rapidly upregulated in separated tops and grafted tops, whereas expression of *DIN6*, *GDH1* and *STP1*, genes whose expression is repressed by sugar (25–27), was rapidly upregulated in separated bottoms and grafted bottoms (Figure 2B, Figure S4). These observations were consistent with sugar accumulation in the grafted top and sugar depletion in the grafted bottom. The expression of these genes returned to levels similar to intact samples by 120 hours and, with the exception of *ApL3*, the grafted samples normalised expression more rapidly than the separated tissues. Genes associated with photosynthesis increase expression in separated bottoms 24 hours after cutting, a common response to starvation (13), but likely too late to affect sugar levels before 24 hours (Figure S4). A transcriptional overlap analysis with RNAs from known glucose-responsive genes (Table S1) revealed a strong overlap with genes differentially expressed by grafting. RNAs from known glucose-induced genes were upregulated in separated tops and grafted tops, whereas transcripts from known glucose-repressed genes were upregulated in separated bottoms and grafted bottoms (Figure 2C, Figure S4). This trend was not observed with genes differentially expressed by mannitol treatment (Figure S4), suggesting the effect was specific to metabolically active sugars. To further investigate this effect, we stained grafted, separated and intact plants with Lugol solution to assay for the presence of starch. Staining above the graft junction increased 48-72 hours after grafting (Figure 2D). By 120 hours, staining was equal on both sides of the graft whereas in separated tops staining became stronger after 72 hours (Figure 2D, Figure S4). We concluded that starch accumulated above the graft junction, but after 72 hours, this asymmetry disappeared. To test whether the accumulation of starch and increased sugar responsiveness could explain the observed transition from asymmetry to symmetry, we compared our datasets to previously published genes that are induced by starvation or are induced by sucrose re-addition (Table S1). In early time points, 20-31% of asymmetrically expressed genes are genes known to respond to sugars compared to 2-5% of symmetrically expressed genes (Table 1). However, at 72 hours, the overlap between asymmetrically expressed genes and sugar responsive genes reduced substantially (Table 1).

**Figure 2.**
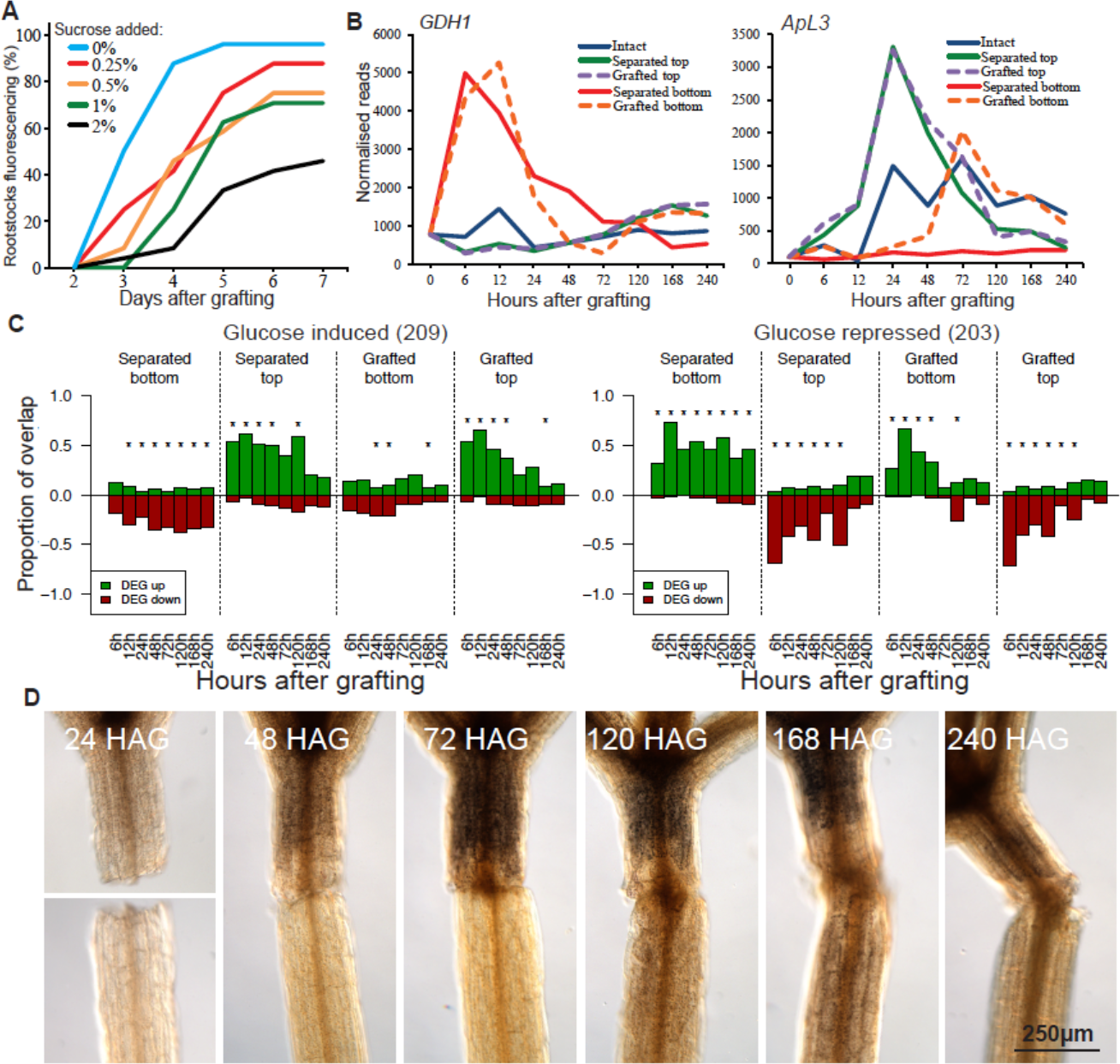
Asymmetric changes in accumulation of sugar-responsive RNAs and of starch occur at the graft junction; equalizing sucrose between the grafted segments reduces and delays phloem reconnection. (A) *pSUC2::GFP*-expressing *Arabidopsis* shoots were grafted to Col-0 wild type roots and GFP movement to the roots was monitored over 7 days for phloem connection in the presence or absence of various concentrations of sucrose. (B) Expression profiles for transcripts of a sugar-repressed gene (*GDH1*), and a sugar-induced gene (*ApL3*) were plotted for intact, separated and grafted samples. (C) Transcriptional overlap between previously published glucose-induced or glucose repressed genes and our dataset. The number in brackets represents the number of glucose-responsive genes identified in the previous dataset, and overlap is presented as a ratio out of 1.0 for differentiation expressed genes (DEG) up- or down-regulated in our dataset relative to intact samples. Asterisks represent a significant difference (p < 0.05) between the ratio of up- and-down- regulated genes in a previously published transcriptome dataset compared to the ratio of all up- and-down- regulated genes in our grafting dataset at a certain time point. (D) Lugol staining of grafted plants at various time points revealed dark brown staining associated with starch accumulation. HAG, hours after grafting.

**Table 1.** Sugar response overlaps with asymmetry. Symmetrically and asymmetrically differentially expressed genes were compared to previously published sugar-responsive genes (28)^1^ and the percent overlap calculated (* p<.05). HAG, hours after grafting.

| HAG | Sugar Induced<sup>1</sup> | Graft Bottom=Top | Overlap | % | Graft Top>Bottom | Overlap | % | Graft Bottom>Top | Overlap | % |
| --- | --- | --- | --- | --- | --- | --- | --- | --- | --- | --- |
| 6 | 2243 | 4988 | 263 | 5 | 6679 | 1403 | 21* | 4971 | 99 | 2 |
| 12 | 2243 | 3473 | 165 | 5 | 7111 | 1444 | 20* | 5657 | 231 | 4 |
| 24 | 2243 | 4135 | 171 | 4 | 6873 | 1414 | 21* | 4942 | 195 | 4 |
| 48 | 2243 | 3689 | 197 | 5 | 6601 | 1241 | 19* | 4915 | 218 | 4 |
| 72 | 2243 | 10421 | 1005 | 10* | 2459 | 228 | 9* | 2019 | 138 | 7 |
| 120 | 2243 | 15012 | 1063 | 7 | 1510 | 220 | 15* | 941 | 115 | 12* |
| HAG | Sugar Repressed<sup>1</sup> | Graft Bottom=Top | Overlap | % | Graft Top>Bottom | Overlap | % | Graft Bottom>Top | Overlap | % |
| 6 | 1998 | 4988 | 107 | 2 | 6679 | 53 | 1 | 4971 | 1563 | 31* |
| 12 | 1998 | 3473 | 68 | 2 | 7111 | 112 | 2 | 5657 | 1526 | 27* |
| 24 | 1998 | 4135 | 88 | 2 | 6873 | 72 | 1 | 4942 | 1525 | 31* |
| 48 | 1998 | 3689 | 113 | 3 | 6601 | 93 | 1 | 4915 | 1427 | 29* |
| 72 | 1998 | 10421 | 530 | 5 | 2459 | 111 | 5 | 2019 | 538 | 27* |
| 120 | 1998 | 15012 | 979 | 7 | 1510 | 84 | 6 | 941 | 210 | 22* |

### Vascular formation and cell division activates on both sides of the graft

Since the grafted bottom samples exhibited a starvation response up to 48 hours after grafting, we reasoned that pathways associated cell division and cell differentiation would be delayed or inhibited. We looked at the expression of markers associated with vascular formation and cell division in the transcriptome datasets. Cambium, phloem and provascular markers activated within 6 hours in grafted top samples, but activation was only delayed 0-24 hours in grafted bottom samples depending on the gene (Figure 1B-C, Figure 3, Figure S5, Figure S6). Expression of phloem markers peaked in both grafted tops and grafted bottoms at 72 hours (Figure 3, Figure S5, Figure S6), the time when phloem reconnections form in grafted *Arabidopsis* (Figure 1, (4, 5)). Notably, the early phloem marker *NAC020* activated before the mid phloem marker *NAC086* which activated before the late phloem marker *NEN4*, consistent with the dynamics of phloem transcriptional activation during primary root development and leaf vascular induction (Figure S5)(29, 30). Certain markers associated with xylem formation, such as *VND7* and *BFN1*, activated early in the grafted top. Other xylem markers, such as *IRX3* and *CESA4*, activated late in grafted samples. By 120 hours after grafting, xylem markers were activated in top and bottom, consistent with when the first xylem strands differentiate at the graft junction (4). Genes associated with cell division were activated by 12 hours in the grafted top and by 24 hours in the grafted bottom (Figure 3, Figure S5). On the other hand, control genes whose expression does not typically vary between tissues and treatments (31) were not differentially expressed in grafted tops or bottoms (Figure S6). The RNAseq expression data appeared to correlate well with transcriptional fluorescent reporters for both activation dynamics and the localisation of expression (Figure 1D-F, Figure S2).

**Figure 3.**
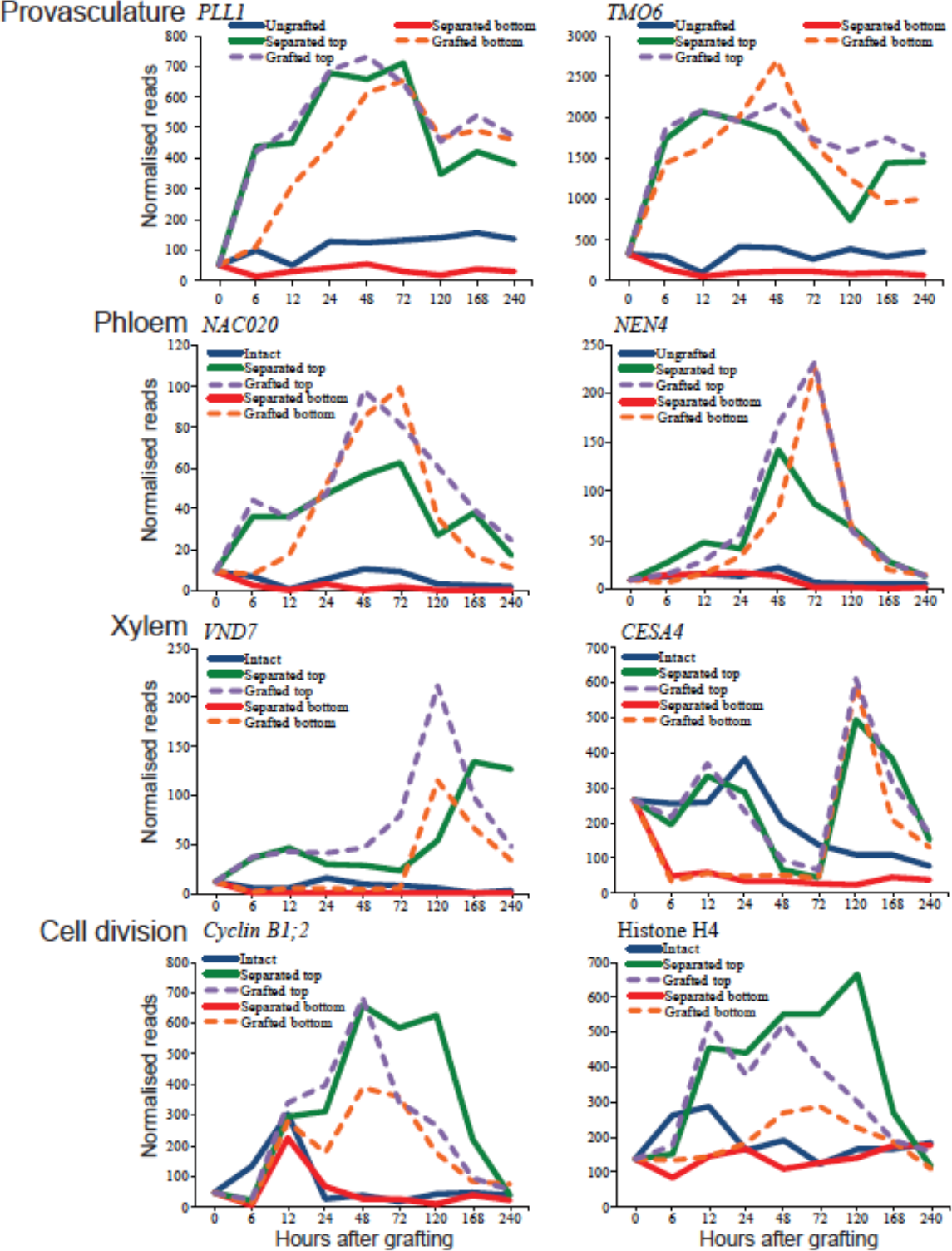
Transcriptional dynamics of genes associated with provasculature, phloem, xylem and cell division. Expression levels were plotted over time for intact, separated and grafted samples.

The similar activation dynamics of vascular differentiation genes in grafted tops and grafted bottoms prompted us to test whether this phenomenon occurred with other known developmental processes. We obtained lists of genes whose expression is associated with various biological processes from previous publications (Table S1) and tested how many of the genes differentially expressed in our transcriptomes overlapped with the previously published lists. Differentially expressed genes in grafted samples and separated tops partially overlapped with those whose expression is associated with phloem, xylem and procambium formation (Figure 4, Figure S7). There was a high overlap between *Arabidopsis* inflorescence stem healing and grafting, as well as between vascular induction in leaf disk cultures and grafting (Figure 4). Various genes expressed in a cell type-specific manner also showed a high transcriptional overlap with grafting, including phloem, endodermis and protoxylem (Figure 4, Figure S7). In nearly all cases, the separated top, grafted top and grafted bottom samples showed similar activation dynamics. The separated bottom samples were exceptional though since gene expression associated with vascular development and cell-specific processes was downregulated (Figure 4, Figure S7). We also compared our datasets to RNAs expressed in longitudinal cross sections of the *Arabidopsis* root (32). There was little overlap between grafted bottoms and sections from the root meristemic zone, whereas overlap existed between grafted tops and the root meristemic zone in early time points, and between grafting and the root maturation zone (Figure S8). Our analysis also revealed that two genes expressed in cambium, *WOX4* and *PXY*, were induced by grafting but the primary root marker *WOX5* and lateral root marker *LBD18* were not (Figure 3, Figure S5, Figure S6).

**Figure 4.**
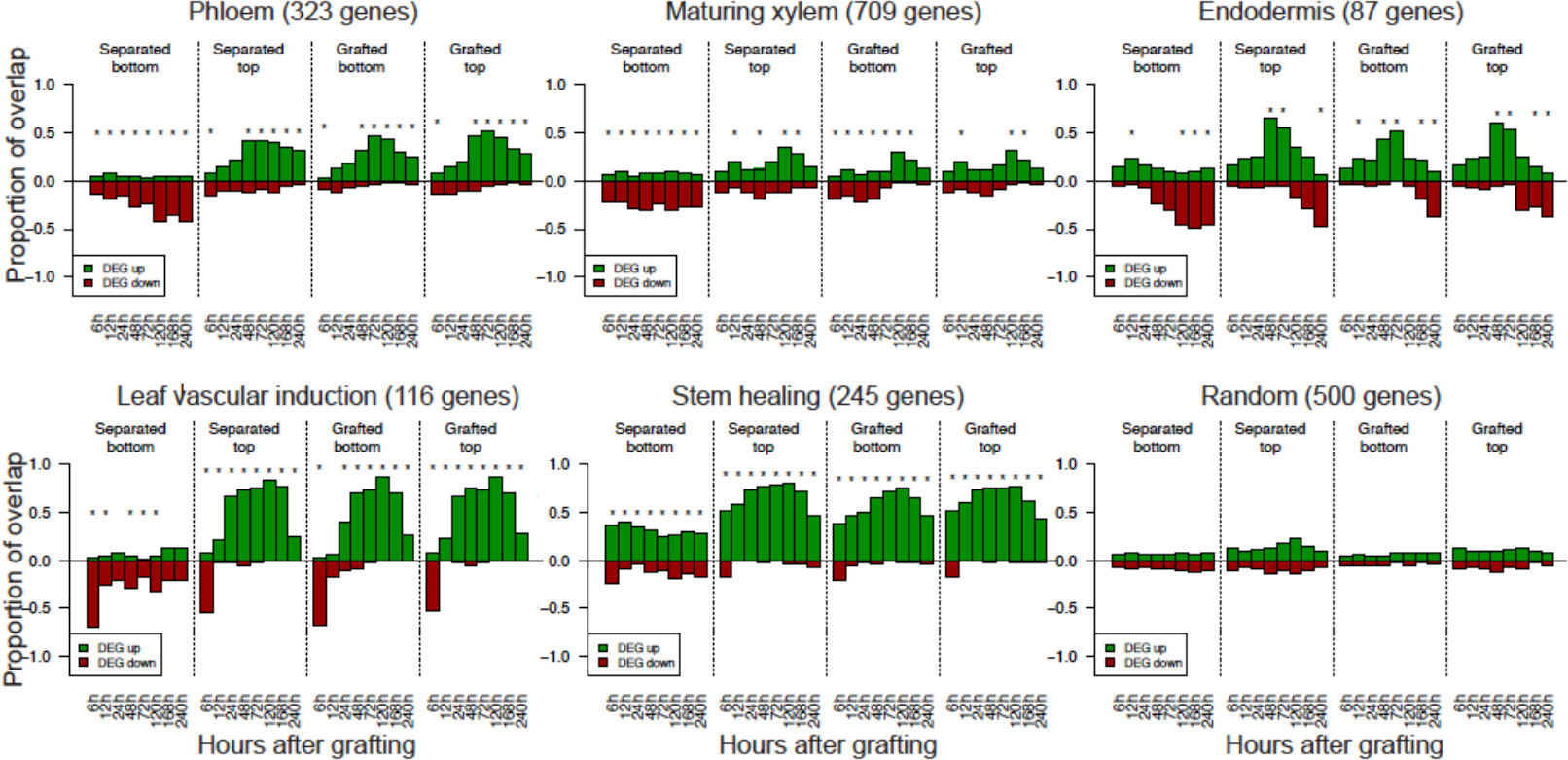
Transcriptional overlap between previously published vascular datasets and the grafting datasets. Genes whose transcripts are associated with various cell types or biological processes were taken from previously published datasets (see Table S1) and compared to the transcriptomic datasets generated here. The number in brackets represents the number of cell type-specific or process-specific genes identified in the previous dataset, and overlap is presented as a ratio out of 1.0 for differentiation expressed genes (DEG) up- or down-regulated in our dataset relative to intact samples. Asterisks represent a significant difference (p < 0.05) between the ratio of up- and down- regulated genes in a previously published transcriptome dataset compared to the ratio of all up- and down- regulated genes in our grafting dataset at a certain time point.

### Auxin response is symmetric at the graft

The rapid activation of vascular markers in the grafted bottoms despite the starvation response promoted us to investigated whether other mobile substances such as phytohormones could play a role in gene activation. We compared lists of genes known to respond to cytokinin, ethylene or methyl jasmonate (33) and found no substantial overlap between these lists and genes differentially expressed by grafting (Figure S9)(Table S1). Abscisic acid-responsive and brassinosteroid-responsive genes showed overlap with genes differentially expressed in our datasets, but this overlap was of a similar magnitude in both separated and grafted datasets suggesting the effect was not specific to grafting (Figure S9). Auxin responsive transcripts were exceptional though, as they showed a substantial overlap with RNAs differentially expressed in our data sets that varied depending on the treatment (Figure 5A-B, Figure S9). Auxin-induced genes were upregulated in separated tops, grafted bottoms and grafted tops whereas they were repressed in separated bottoms (Figure 8). Auxin responsive genes such as *IAA1* and *IAA2* (34) were induced to similar levels in grafted tops and grafted bottoms at 24 hours. To further investigate whether auxin response was uniform between grafted tops and grafted bottoms, we grafted the auxin-responsive fluorescent reporter *p35S:DII-Venus* whose fluorescent protein is degraded in the presence of auxin (35). DII-Venus fluoresced in the separated bottoms but did not fluoresce in grafted bottoms 14 hours after cutting (Figure 5C) indicating separated bottoms had a low level of auxin response but grafted tops, grafted bottoms and separated tops had a high level of auxin response.

**Figure 5.**
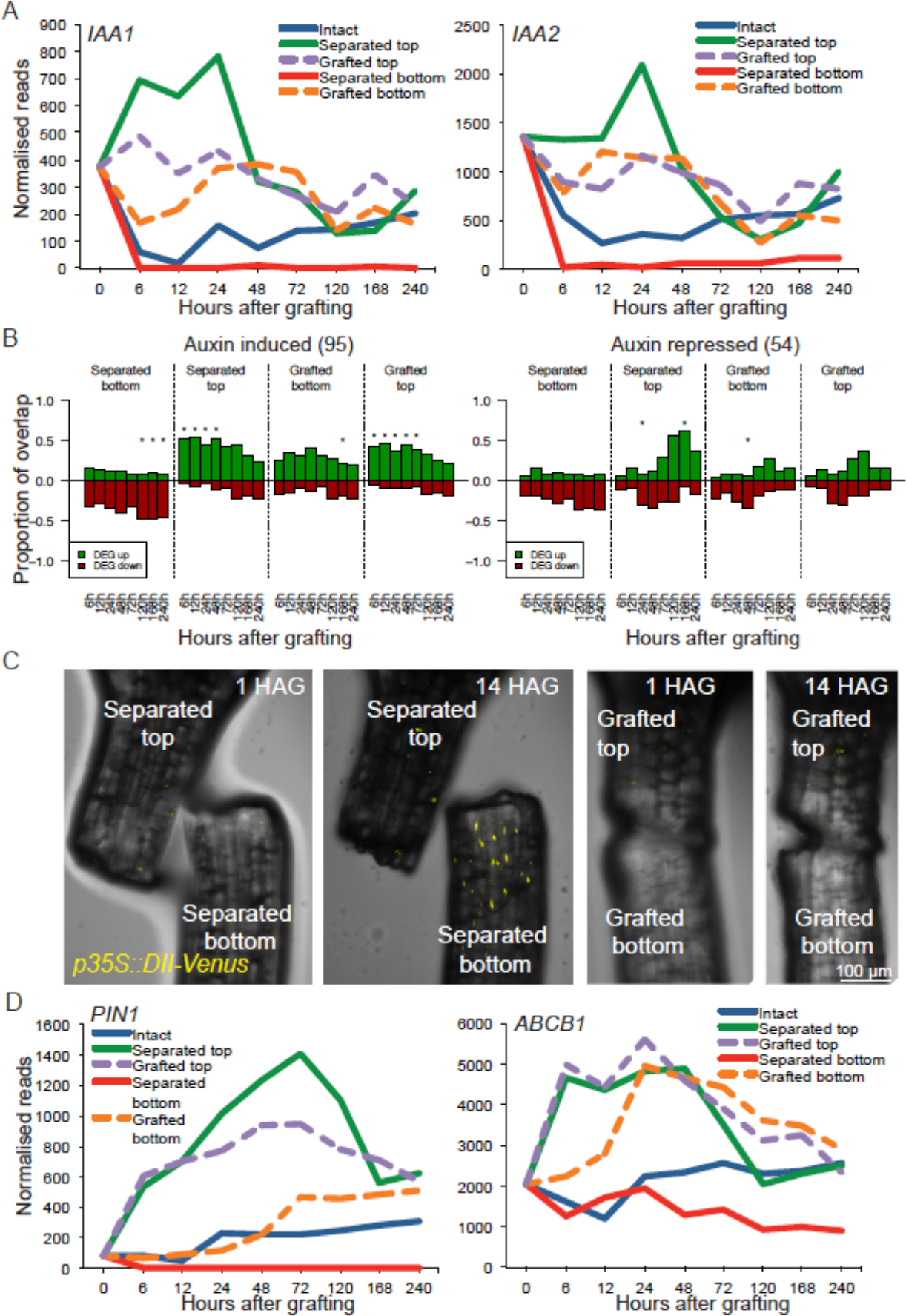
Auxin response is symmetric at the graft junction. (A,D) Expression profiles for various auxin-responsive genes (*IAA1*, *IAA2*) or auxin transporter genes (*PIN1* and *ABCB1* genes) were plotted for intact, separated and grafted samples. (B) Overlap between previously published auxin-induced or auxin-repressed RNAs and our dataset. The number in brackets represents the number of auxin-responsive genes identified in the previous dataset, and overlap is presented as a ratio out of 1.0 for differentiation expressed genes (DEG) up- or down-regulated in our dataset relative to intact samples. Asterisks represent a significant difference (p < 0.05) between the ratio of up- and down- regulated genes in a previously published transcriptome dataset compared to the ratio of all up- and down- regulated genes in our grafting dataset at a certain time point. (C) Grafted and separated plants expressing the auxin responsive *p35S::DII-Venus* transgene that is degraded in the presence of auxin reveal a reduction of auxin response in cut bottoms, but not in grafted bottoms. HAG, hours after grafting.

### Tissue fusion imparts a unique physiological response that differs from tissue separation

We hypothesized that the symmetric auxin response and asymmetric sugar response at the graft junction could allow a unique transcriptional response since neither separated plants nor intact plants had similar response dynamics to sugars and auxin as measured by genome-wide gene response, starch accumulation and *p35S::DII-Venus* expression (Figure 2, Figure 5). To uncover protein-coding genes differentially expressed specifically by grafting, we performed an empirical Bayesian analysis (36) where at each time point we computed likelihoods for each gene for each possible pattern of differential expression in the intact, grafted top, grafted bottom, separated top and separated bottom tissues (Figure 6A). These likelihoods were used to define a similarity score between pairs of genes, which was used to cluster genes with similar patterns of gene expression across the five tissues (37). Clusters were then identified by the predominant pattern of gene expression observed within a cluster. This analysis produced 113 clusters containing at least 10 RNAs at a single time point (Table S2; Dataset S1). Approximately 6000 genes were differentially expressed in at least one tissue whereas between 1000 and 4000 genes were not differentially expressed (Figure 6B).

**Figure 6.**
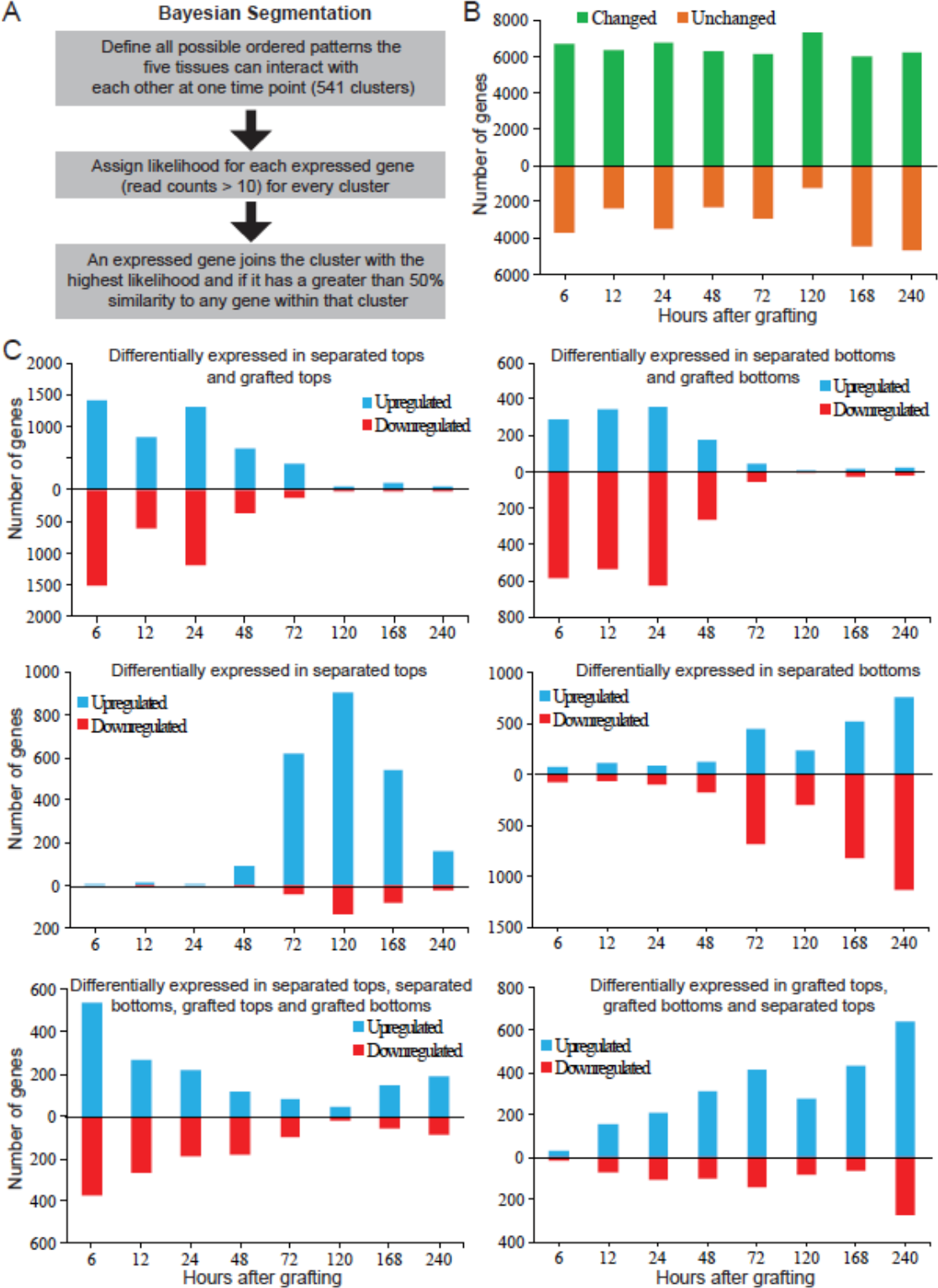
Clustering the transcriptome at each time point, based on likelihoods of all possible patterns of differential expression. (A) Cartoon depicting the Bayesian segmentation. (B) Analysis of differential behaviour produced 113 categories containing at least 10 genes whose expression was expressed in a specific differential pattern in at least one time point (see Table S2). One group is comprised of genes whose transcript levels are not substantially changed in the five tissues (unchanged) whereas the other group is comprised of the sum of the other 112 groups (genes whose transcript levels changed after treatment in a least one tissue) over the time points tested. (C) Major categories in the segmentation revealed RNAs whose levels changed in all the treatments listed relative to intact samples. Note that a gene can be represented only in one category for a given time point, that category in which the transcript level changes best fit the category.

To simplify the analysis, we considered clusters where gene expression was grouped into patterns consisting of one comparison. At early time points, the cluster containing genes differentially expressed in grafted tops, grafted bottoms, separated tops and separated bottoms had high numbers that decreased with time and could represent an general wound response (Figure 6C). Similarly, clusters containing separated tops and grafted tops and clusters containing separated bottoms and grafted bottoms initially had high numbers that decreased with time. This observation indicated the grafted top was initially transcriptionally similar to the separated top, whereas the grafted bottom was initially transcriptionally similar to the separated bottom. After the 48-hour time point, clusters containing genes only differentially expressed in separated tops or only differentially expressed in separated bottoms increased in numbers, suggesting these tissues gained a unique pattern of gene expression. The cluster containing genes differentially expressed in grafted tops and grafted bottoms increase in numbers throughout the healing process (Figure 6C). We searched for cluster categories that contained genes only differentially expressed by grafting and found very few genes downregulated by grafting or upregulated only in the grafted top (Figure 7A). Instead, there were clusters that contained several hundred differentially expressed genes present either in the grafted bottom only, or present in both grafted bottom and grafted top (Figure 7A). Genes whose expression changed only in the grafted bottom sample were prevalent early during grafting and were most common at 48 hours, whereas genes activated in both top and bottom became prevalent at 48 hours and were most common at 120 hours (Figure 7A). We performed a gene ontology (GO) analysis on the genes whose expression was grafting-specific and found that genes coding for RNAs differentially expressed only in the grafted bottom sample were enriched in the immune response and chitin response biological process categories (Table S3). Previously published chitin-induced RNAs had a high proportion of overlap with differentially expressed graft bottom-specific genes (Figure 7E, Figure S10). Grafting-specific RNAs expressed symmetrically, in both the grafted top and grafted bottom, were enriched in vascular-related biological processes (Table S3). Previously published phloem-enriched, endodermal-enriched, vascular-induction and stem-wounding associated RNAs had a high proportion of overlap with differentially expressed graft-specific genes (Figure 7C-D, Figure S10). Since few genes were grafting-specific and grafted tissues were initially transcriptionally quite similar to separated tissues (Figure 6C, Figure 7A), we reasoned that tissues separated for short periods (<48 hours) could be grafted with similar reconnection dynamics as tissues that had been grafted immediately. To test this hypothesis, plants were cut at day 0, and grafted at 1-5 days after separation. Separated tissues did not speed up vascular reconnection, and instead, it always took three days from the point of tissue attachment for vascular connections to form (Figure S10). Furthermore, the shoot lost competence to graft 2-3 days after separation whereas the root remained competent to graft up to 5 days after separation (Figure S10). Together, it appears the grafted shoot and root have a unique physiological response that differ from separated plants and that tissue attachment is required to activate graft formation and tissue fusion.

**Figure 7.**
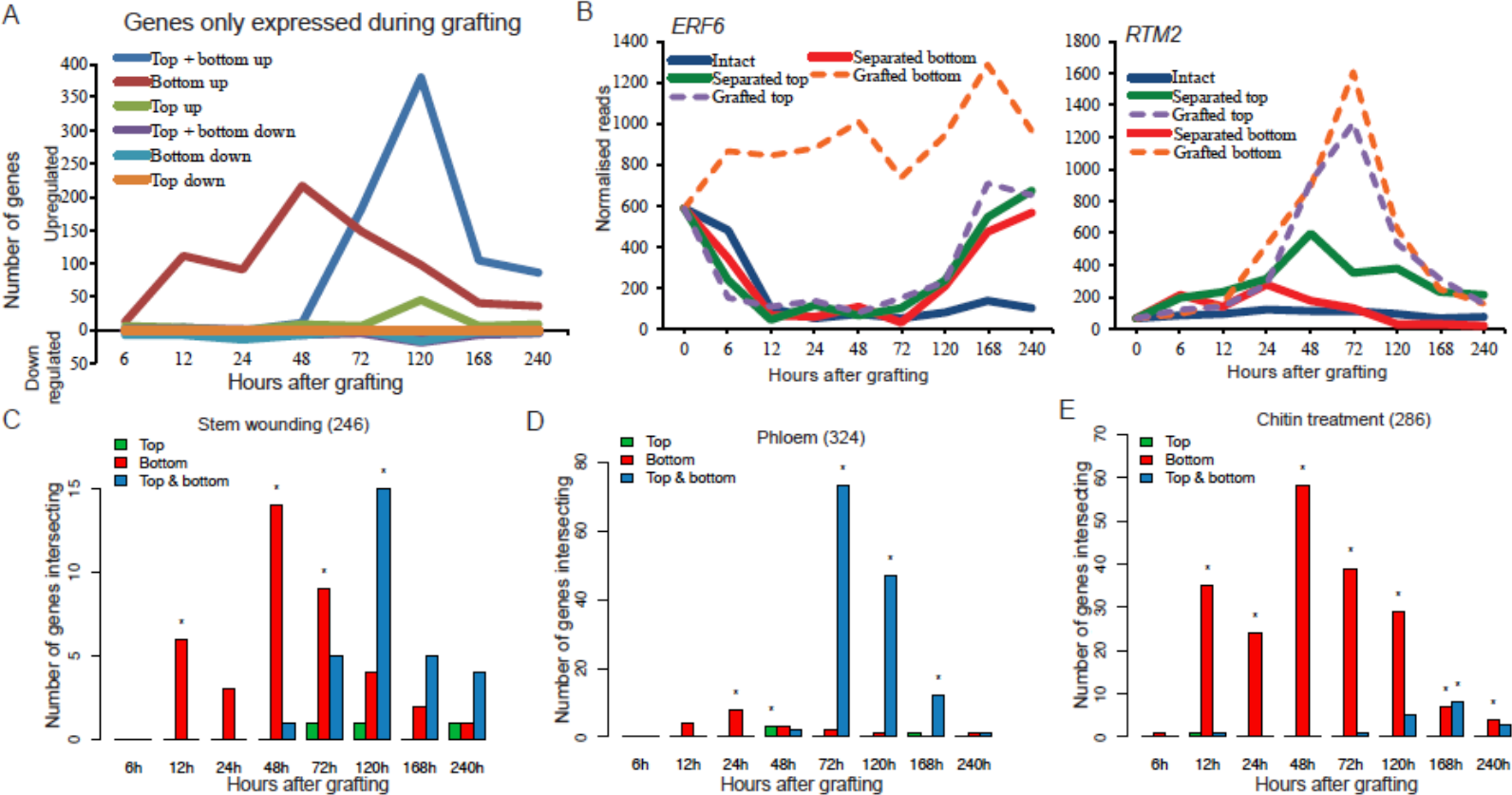
A subset of genes is differentially expressed only during graft formation when compared to 0 hour grafts, or intact or separated tissues. (A) Certain genes were only differentially expressed in grafted tops, grafted bottoms or both in grafted tops and grafted bottoms. (B) Expression profiles for a graft bottom-specific (*ERF6*) or a graft top and bottom differentially regulated gene (*RTM2*) were plotted for intact, separated and grafted samples. (C) Grafting-specific genes are also expressed in other processes of tissue fusion, such as stem healing. Here, 246 previously published RNAs whose differential expression is associated with stem healing were compared to our dataset to assess transcriptional overlap with the grafting-specific genes. Asterisks represent a significant high overlap (p <0.05) of previously published genesets that are also differentially expressed in the grafted samples at a certain time point. (D-E) Genes differentially expressed in grafted tops and grafted bottoms show high overlap to 324 previously published genes whose transcripts are associated with phloem, whereas genes expressed in grafted bottoms show high overlap to 286 previously published genes whose encoded RNAs are associated with chitin treatment. Asterisks represent a significant high overlap (p <0.05) of previously published genes that are also differentially expressed in the grafted samples at a certain time point.

## DISCUSSION

To better understand how plants graft, we analysed in depth an RNA deep sequencing dataset that spatially and temporally distinguished genes activated by cutting followed by tissue attachment or continuous tissue separation. Cutting promoted a similar wound response in both grafted and separated tissues, however, by 72 hours after cutting the grafted and separated tissues became transcriptionally dissimilar (Figure 6C) indicating that tissue fusion was mechanistically different from healing an unattached cut surface. During graft formation, tissues had a very high transcriptional overlap with genes differentially expressed by inflorescence stem healing and by vascular induction in leaves (Figure 4, Figure S7) (9, 29) suggesting grafting is closely related to tissue reunion and vascular formation processes. Graft formation had little transcriptional overlap with lateral root formation (Figure S6)(32) and appeared to follow a pathway similar to secondary root growth (Figure S6, Figure S8) since cambium markers *WOX4* and *PXY* that are specific to secondary growth were expressed during grafting (38)(Figure S5). Grafted tops initially showed a short-lasting and small transcriptional overlap with genes expressed during primary root formation which may be related to the accumulation of substances activating adventitious root formation, a common response in failed grafts or in cut shoots (Figure S4C). Thus, we conclude that grafting proceeds via a pathway involving secondary growth with radial meristems activating in the mature cambium to heal the wound. Vascular formation genes including those specifying cambium and phloem were activated early, followed by an activation of cell division genes, suggesting that the start of cellular differentiation preceded activation of cell division. Xylem identity genes showed an early and a late activation peak (Figure 3, Figure S5). There is no visible xylem differentiation at the graft junction during the first peak of expression (4) and these genes might be suppressed by phloem differentiation genes such as *APL* and *CLE41* that activate early after grafting and are known to suppress protoxylem formation (39, 40)(Figure S6)(Table S4). Alternatively, the first peak could represent programmed-cell death that does not lead to xylem differentiation. The second expression peak of xylem-expressed genes at 120 hours occurred after the differentiation of functional phloem and coincided with the differentiation of xylem strands at the graft junction (4). Previous studies highlighted the importance of callus and pericycle cells during regeneration (16, 41), but we see little evidence that genes expressed in the pericycle or during callus formation have high transcriptional overlap with genes differentially expressed by grafting (Figure S4). Expression profiles for all protein-coding genes can be found in table S4.

We observe only a slight delay in phloem, cambium and cell division activation below the graft junction compared to above it (Figure 3, Figure S5, Figure S6) whereas several genes associated with vascular formation, such as *HCA2*(42) and *TMO6*(43), activated equally in both grafted top and grafted bottom at 6 hours after grafting (Figure 1D-G, Figure 3). These data indicate that, at least transcriptionally, the grafted root rapidly responded to the presence of the grafted shoot and this response was independent of functional vascular connections. This response was not present in separated roots, indicating that attachment was key for recognition. Sugars are known activators of cell division and cell elongation (13) and in our datasets, a large proportion of asymmetrically expressed genes are genes that are sugar-responsive. However, sugars are transported in the phloem (12) that is severed upon grafting whereas the grafted root exhibited a sugar starvation response and showed similar sugar-response dynamics as the separated root. Instead, we infer that not sugar but auxin, or some other molecular that is transported in the absence of vascular connections, could be the signal in the grafted bottom that activates *HCA2*, *TMO6* as well as cell division, phloem- and cambium-related genes. Alternatively, separation might cause a build-up of molecules from the root that suppress vascular induction genes such as *HCA2* and *TMO6*.

Given auxin’s role in vascular patterning and formation ((38), it is a strong candidate for the activating signal. Auxin response was largely symmetric, particularly from 12 hours after grafting (Figure 6) consistent with previous findings that the auxin-inducible *DR5* and *ANAC071* genes are symmetrically expressed around the graft junction (4–6, 44). One idea is that auxin transport was not substantially interrupted by the grafting process, and instead, where opposing tissues adhered, auxin moved regardless of vascular connections since auxin is transported from cell to cell through the apoplast (8). The genes encoding auxin efflux proteins PIN1 and ABCB1 were transcriptionally activated above the graft junction (Figure 5D), similar to putative *Pisum sativum* PIN1 protein accumulating above a cut stem prior to vascular reconnection (45), and could reflect a role for these proteins in exporting auxin across the cut. Consistent with these observations, adding an auxin transport inhibitor to grafted *Arabidopsis* shoots prevented the expression of grafting-induced genes below the graft junction (6). Since auxin is fundamental for patterning vasculature (38), it is plausible that auxin movement across the graft occurs prior to sugar movement since the flow of auxin could assist with reconnecting the phloem. Although auxin response was symmetric, our previous work demonstrated that the auxin response factors *ALF4* and *AXR1* affected grafting only below the graft junction (4). Mutating *ALF4* below the graft junction more strongly reduced auxin response than mutating *ALF4* above the junction (4). Thus, proteins such as ALF4 or AXR1 might enhance or promote rootstock-specific auxin response and vascular regeneration that could be particularly important when there is incomplete attachment, cellular damage or inefficient transport. All higher plants transport auxin from shoot to root, yet not all plant species can be successfully grafted (3) so the response to auxin rather than the transport itself may be a determining factor in the ability to graft. A role for sugars is not completely ruled out though, since the magnitude of differential expression was often lower in the grafted bottom and exogenous sugar reduced grafting efficiency (Figure 2A). Pericycle cells require auxin to divide and the addition of sugars enhances the rate of divisions (46), suggesting that the presence of sugars in the grafted top enhanced cell division and differentiation. Another hypothesis that warrants testing is whether graft healing activates via a mechanical signal provided by the physical presence of the opposing tissues.

Our analyses identified two groups of genes whose expression changes were unique to graft formation in our experiments (Figure 7). One group activated shortly after grafting below the graft junction and was enriched in immune-responsive and chitin-responsive genes (Table S3, Figure 7, Figure S10). The breakdown products of cell walls are potent elicitors of defence responses (47), so it is possible that the grafted bottom upregulates pathways specific to wound damage response. This group was not upregulated in separated bottoms though, so the unique physiological state of the grafted root, indicated by the presence of auxin response but the absence of sugar response, could have promoted their upregulation. The second group activated both above and below the graft junction and became highly expressed later during graft formation (Figure 7). This group was enriched in RNAs associated with vascular development (Table S3, Figure 7) and we suggest that the products of these genes are involved in the vascular reconnection processes between the two tissues. Despite many transcriptional similarities between separated tops and grafted tissues, tissues had to be attached for at least three days for phloem connections to form, regardless of when cutting occurred (Figure S10). Thus, it appears that RNAs expressed in the separated top or separated bottom are insufficient to drive graft formation. Instead attachment and the genes activated by this recognition process including grafting-specific RNA changes (Figure 7A) or RNAs expressed in the grafted bottom, grafted top and separated top (Figure 6C) are those that contribute to a response that distinguishes attached from separated plant tissues. Future work should focus on these RNAs to identify the pathways required for grafting that could be modified to improve graft formation, wound healing and vascular regeneration. Likewise, the rapid transcriptional changes below the graft indicate a recognition system that promotes tissue regeneration. Identifying the exogenous cue that triggers recognition and understanding how it is perceived should be priorities, as should understanding whether this phenomenon applies more broadly to inter-tissue communication, tissue regeneration or tissue fusion events, such as parasitic plant infections (48), epidermal fusions (49, 50) or petal fusions (51).

## METHODS

### Plant material and microscopy

*Arabidopsis thaliana* accession Columbia was used throughout. The *p35S::GFP-ER* (52), *pSUC2::GFP* (53), *pUBQ10::PM-tdTomato* (54), *pARR5::GFP* (55), *pANT::H2B-YFP* (56), *pLOG4::n3GFP* (57), *pCASP1::NLS-GFP* (58), *p35S::DII-Venus* (35) lines have been previously published. For the construction of *pHCA2::RFP*, a 2.9kb 5’ upstream region of the *HCA2* gene was cloned into pDONRp4-p1R donor vector, and recombined with tagRFPer into a destination vector by the Multisite Gateway system (59). *Arabidopsis thaliana* micrografting and grafting assays were performed according to previously published protocols (60, 61). Fluorescent images were taken on a Zeiss LSM-700 or LSM-780 confocal microscope. Black and white fluorescent images of graft junctions were taken on a Zeiss V12 dissecting microscope. FIJI software (Fiji.sc) was used to process images. A micro-extensometer was used for all breaking force graft measurements according to a previously published protocol (62).

### RNAseq sample and library preparation

Grafted wild type *Arabidopsis thaliana* accession Col-0 were harvested at the respective time points and care taken to separate grafts by gently pulling plants apart. Approximately 0.5mm of tissue was taken above or below each cut site and kept separate. Intact plants had 1mm of tissue taken in a similar location on the hypocotyl as separated or grafted plants. Grafted, separated or intact tissues were pooled into groups of approximately 24 tissues. Tissues were ground using a microcentrifuge pestle frozen in liquid nitrogen. RNA was extracted using an RNeasy Kit (Qiagen, UK) following the manufacturer’s instructions. 90-100ng of RNA was used to prepare RNAseq libraries using the TruSeq^®^ Stranded mRNA LT kit (Illumina, UK) according to the manufacturer’s instructions. The final PCR was for 15 cycles and 11-12 barcoded samples were randomly mixed to make a total of 7 mixes for 7 flow lanes, one mix per lane. Samples were sequenced on the HiSeq 4000 platform (Illumina, UK) with Paired End 75bp transcriptome sequencing (BGI Tech Solutions, Shenzhen, China).

### Iodine staining

*Arabidopsis* seedlings were placed in a fixation solution (3.7% formaldehyde, 50% ethanol, 5% acetic acid) for 1 hour at room temperature, then transferred to 70% ethanol for 10 minutes. Afterwards, plants were transferred to 96% ethanol and stored at −20°C for up to a week. Samples were rehydrated in 50% ethanol for 1 hour at room temperature, transferred to distilled water for 30 minutes, then stained for 10 minutes in Lugol solution (Sigma) at room temperature. Plants were rinsed with water and mounted on microscope slides. Images were taken on a Zeiss Axioimager.M2 microscope.

### Bioinformatic analyses

The reads acquired through high-throughput sequencing were quality trimmed with sickle (63) and aligned using the eXpress tool to protein-coding gene sequences acquired from TAIR10 using Bowtie2. Library scaling factors were inferred from the sum of the number of reads assigned to the genes in the lowest seventy-five percentiles of expressed genes for each library (64). Analyses of the data were carried out using the R package baySeq (36)and clustering based on the posterior probabilities acquired from this package. The gene ontology analysis (GO) enrichment analysis on grafting-specific genes was done with a customized R script using the package GOstats (65).

## ACKNOWLEDGEMENTS

We thank Niko Geldner, Dolf Weijers, Paul Tarr, Yka Helariutta, Ruth Stadler and The Arabidopsis Information Resource for providing seeds. Funding for this work was provided by Gatsby Charitable Trust grants GAT3272/C and GAT3273-PR1, by a Knut and Alice Wallenberg Academy Fellowship KAW2016.0274 (to C.W.M), and by the Howard Hughes Medical Institute and Gordon and Betty Moore Foundation grant GBMF3406 (to E.M.M.)

## SUPPORTING INFORMATION

### SI Materials and Methods

#### Plant material and grafting

*Arabidopsis thaliana* accession Columbia was used throughout. The *p35S::GFP-ER* (52), *pSUC2::GFP* (53), *pUBQ10::PM-tdTomato* (54), *pARR5::GFP* (55), *pANT::H2B-YFP* (56), *pLOG4::n3GFP* (57), *pCASP1::NLS-GFP* (58), *p35S::DII-Venus* (35) lines have been previously published. For the construction of *pHCA2::RFP*, a 2.9kb 5’ upstream region of the *HCA2* gene was cloned into pDONRp4-p1R donor vector, and recombined with tagRFPer into a destination vector by the Multisite Gateway system (59). The following primers were used for *pHCA2* cloning: attB4_HCA2(‒)2958 GGGACAACTTTGTATAGAAAAGTTGtcgatacgcgggacagatatac attB1_HCA2ProEnd ggggACTGCTTTTTTGTACAAACTTGttttgtgttctgtatgtttg. *Arabidopsis thaliana* micrografting was performed according to a previously published protocol (60). Briefly, seven day old *Arabidopsis* seedlings were grown vertically on Murashige and Skoog (MS) medium + 1% bacto agar (pH5.7; no sucrose) in short day conditions (8 hours of 80-100 µmol m^−2^ s^−1^ light) at 20°C. Seedlings were placed on one layer of 2.5x4cm sterile Hybond N membrane (GE Healthcare) on top of two 8.5cm circles of sterile 3 Chr Whatman paper (Scientific Laboratory Supplies) moisten with sterile distilled water in a 9cm petri dish. In a laminar flow hood using a dissecting microscope, one cotyledon was removed and a transvers cut through the hypocotyl was made with a vascular dissecting knife (Ultra Fine Micro Knife; Fine Science Tools). Grafts were assembled by aligning the two cut halves and joining them together, after which, the petri dishes was sealed with parafilm and placed vertically under short day conditions at 20°C. For grafting on a microscope coverslip to image the graft junction, a 10cm square Petri dish modified by gluing a microscope coverslip in place of a section of plastic from the back. On top of the microscope cover slip was placed a 2.5x4 cm rectangle of Hybond N membrane. At the edges and base of the Petri dish three 3 × 8cm strips of Whatman paper were placed. Sterile water moistened both Whatman paper and Hybond N. After which, roots were placed on the Hybond N membrane and hypocotyls on the coverslip. Grafting then proceeded as above. Graft junctions were imaged through the coverslip with a Plan-Apochromat 20X/0.8 objective on a Zeiss LSM-700 or LSM-780 confocal microscope.

#### Fluorescent assays and microscopy

To test the effect of sugars on grafting, *Arabidopsis thaliana* Col-0 plants were grown on 1/2MS+ 1% bacto agar (pH5.7; no sucrose) for seven days in short day conditions. Wild type roots were grafted to scions expressing *pSUC2::GFP* using the protocol described above but either water or water containing 0.25%, 0.5% 1% or 2% sucrose was added to the grafted plates. Roots were observed for fluorescence 2-7 days after grafting with a Zeiss V12 dissecting microscope equipped with a GFP filter. Roots were scored daily and the same plants were observed during the 7-day assay as previously described (61).

Fluorescent images were taken on a Zeiss LSM-700 or LSM-780 confocal microscope with a Zeiss Plan-Apochromat 20X/0.8 dry objective. A 488nm argon laser (Zeiss 780) or 488nm solid-state laser (Zeiss 700) was used for excitation of GFP and YFP. A 561nm solid-state laser was used for excitation of the tdTomato fluorescent protein. A T-PMT detector obtained bright-field transmitted light. Black and white fluorescent images of graft junctions were taken on a Zeiss V12 dissecting microscope fitted with a Hamamatsu EM-CCD camera and RFP and YFP filters. FIJI software (Fiji.sc) was used to process images. Image contrast and brightness were adjusted for controls and samples equally. For longitudinal images of the graft junction, z-stack projections are shown and made with the average intensity function in FIJI from stacks containing the hypocotyl vascular tissues, mesophyll and epidermis.

#### RNAseq sample and library preparation

Wild type *Arabidopsis thaliana* accession Col-0 were grafted as above taking care to switch shoot and root between different plants. All grafting and cutting was performed in the morning to minimize circadian effects. For the 0 hour time points, plants were transferred from 1/2MS plates to the grafting plates and immediately harvested. Only intact plants were harvested at the 0 hour time since at this point, there would be insufficient time to reasonably expect the separated or grafted samples to be transcriptionally different (the time between cutting and freezing is less than two minutes). For all other time points, after cutting plants were left on grafting plates for the respective amount of time. Tissues were harvested and care taken to separate grafts by gently pulling plants apart. Approximately 0.5mm of tissue was taken above or below each cut site and kept separate. Intact plants had 1mm of tissue taken in a similar location on the hypocotyl as separated or grafted plants. Grafted, separated or intact tissues were pooled into groups of approximately 24 tissues (1 plates with 24 plants) which were immediately placed in 96% ethanol on dry ice. After harvesting, microcentrifuge tubes were briefly centrifuged and the ethanol removed before storing at −80°C. Plants were grafted over two months to get sufficient material.

Tissues were ground in the microcentrifuge tube using a microcentrifuge pestle frozen in liquid nitrogen. RNA was extracted using an RNeasy Kit (Qiagen, UK) following the manufacturer’s instructions including on column DNase digestion. RNA was eluted from the column with 50ul of sterile water. Quality and quantity of RNA was checked using an Agilent 2200 TapeStation and High Sensitivity (HS) RNA screentapes (Agilent, UK). After RNA extraction, two to four biological replicates were combined (50-100 plants) to get sufficient RNA. 90-100ng of RNA was used to prepare RNAseq libraries using the TruSeq^®^ Stranded mRNA LT kit (Illumina, UK) according to the manufacturer’s instructions. The final PCR was for 15 cycles and samples were resuspended in 23ul of distilled water. Quantity and quality of DNA libraries was checked on the Agilent 2200 TapeStation using D1000 screentapes (Agilent, UK). Each sample had two libraries prepared from grafted tissues or separated tissues at different times so that independent biological replicates were made. Samples were diluted to 10nM and 11-12 barcoded samples randomly mixed to make a total of 7 mixes for 7 flow lanes, one mix per lane. Samples were sequenced on the HiSeq 4000 platform (Illumina, UK) with Paired End 75bp transcriptome sequencing (BGI Tech Solutions, Shenzhen, China).

#### Iodine staining

*Arabidopsis* seedlings were placed in a fixation solution (3.7% formaldehyde, 50% ethanol, 5% acetic acid) for 1 hour at room temperature, then transferred to 70% ethanol for 10 minutes. Afterwards, plants were transferred to 96% ethanol and stored at −20°C for up to a week. Samples were rehydrated in 50% ethanol for 1 hour at room temperature, then transferred to distilled water for 30 minutes. Samples were then transferred to a solution of Lugol solution (Sigma) and stained for 10 minutes at room temperature. Plants were rinsed with water, then mounted on microscope slides. Images were taken on a Zeiss Axioimager.M2 microscope with a PlanApochromat 20x objective and SPOT Flex camera (Imsol, UK).

#### Pairwise and Bayseq analyses

The reads acquired through high-throughput sequencing were quality trimmed with sickle (63) to increase the read quality before mapping. Reads were aligned to protein-coding gene sequences acquired from TAIR10 using Bowtie2. Read assignment was performed using the eXpress tool, which also provided effective gene lengths for use in normalisation. Library scaling factors were inferred from the sum of the number of reads assigned to the genes in the lowest seventy-five percentiles of expressed genes for each library (64).

Analyses of the data were carried out using the R package baySeq (36)and clustering based on the posterior probabilities acquired from this package. For each timepoint, all possible patterns of differential expression between the graft types were considered, where a ‘pattern’ defines similarity and difference between different experimental conditions. For example,

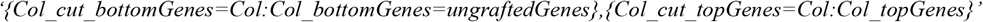

defines a pattern in which gene expression is equivalent in the separated bottoms, the grafted bottoms and the intact plant, but different to the equivalently expressed separated top and grafted top. The total number of possible patterns for five experimental conditions (as in this analysis) is fifty-two.

For a given timepoint, posterior likelihoods on the likelihood of each pattern of expression are calculated for every gene with greater than ten reads across all experimental conditions. The patterns were then modified to include orderings (denoted by < or >), for example, the pattern described would lead to the ordered pattern

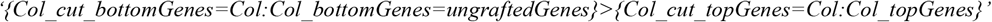

in which gene expression is equivalent in the separated bottoms, the grafted bottoms and the intact plant and greater than the equivalently expressed separated top and grafted top. In total, 541 ordered patterns exist in this data set. Posterior likelihoods for an ordered pattern were assigned to that of the unordered pattern for genes in which the (normalised) mean expressions within the equivalently expressed groups conformed to the ordering, and to zero otherwise.

Based on the posterior likelihoods for the ordered patterns, a similarity score *s*_*ij*_ was established between two genes *i* and *j* as the sum over the products of their likelihoods of each ordered pattern. A single-link agglomerative clustering of genes, in which a gene will join a cluster if it has a greater than 50% similarity to any gene within that cluster was then performed based on these similarity scores. We label each cluster according to the predominant ordered pattern with high likelihood amongst the genes that comprise it. The change in size of these clusters over time is shown for the major clusterings in Figure 6.

We can also find likelihoods on comparisons between pairs of experimental conditions by summing the likelihoods over combinations of patterns. Figure 1B shows the number of genes identified at each time point in a pairwise analysis between the grafted top and grafted bottom samples. The likelihood of symmetric expression (i.e., expression which is equivalent across the graft junction) is calculated as the sum of the likelihoods of all patterns in which the grafted top and grafted bottom samples are equivalent. Conversely, asymmetric expression is calculated as the sum of the likelihoods of all patterns in which the grafted top and grafted bottom samples are not equivalent. Additional sets can be formed by considering the ordering of the grafted top and grafted bottoms samples. Sets of genes are identified at each time point with an FDR of less than 0.05 and a likelihood of symmetric/asymmetric expression greater than 50%. Genes in this analysis were only included if they were differentially expressed relative to intact samples.

#### Gene overlap analyses of up and down regulated genes

To measure if the ratio of up-and down-regulated genes from a previously published dataset is significantly different to the ratio of up-and down-regulated genes in our grafting dataset we only took into account genes that are differentially expressed at a certain time point. A gene was called differentially expressed at a certain time point if the marginal likelihood, calculated by BaySeq, was greater than 0.9 and if the absolute log_2_-foldchange was greater than 1. Hence, we only consider genes that are significantly two-fold up- or down- regulated. This definition of differentially expressed genes was also used to filter the published datasets according to the expression values in our transcriptome dataset. Hence, some genes were filtered out from the original published datasets because they did not show a significant up- or down- regulation during a certain time point in our expression data based on our criteria. The histograms (Figure 2,4, 5, S4, S7, S8, S9) show the relative number of up- and down- regulated genes from the published datasets during a certain time point and a certain condition (separated top, separated bottom, grafted top, grafted bottom) based on the number of genes in the published dataset after filtering. To calculate the significance of the difference of the ratios between the published DEGs and all up- and down- regulated genes, we used a two-sided Fisher’s exact test. To correct for multiple testing we used the Benjamini-Yekutieli (BY) correction method. Hence, the asterisks in the barplots highlight that the corrected p-value is below 0.05.

#### Dealing with probe ids from microarray datasets

Due to the fact that some published datasets only used probe ids instead of gene ids to represent their differentially expressed genes we first had to match these probe ids to their corresponding gene ids. This step was done with the R package biomartr (66). If one probe id matched more than one gene id we used all the corresponding gene ids and tested afterwards if these genes were actually differentially expressed in our dataset. In some cases, one probe id was represented by more than one gene id. Hence, some gene sets contained slightly more gene ids than published probe ids. In contrast, some probe ids did not match to currently existing gene ids. Hence, some gene sets contained slightly fewer gene ids than published probe ids.

#### Gene overlap analyses of gene sets involved in graft formation

For this analysis we used the previously calculated gene sets from baySeq of differentially expressed genes clustered for each time point into grafted bottom, grafted top, and grafted bottom and top and calculated their overlap to previously published data sets. The significance of the number of overlapping genes between the grafted samples and the published datasets was determined by a one-sided Fisher’s exact test, to prove if the overlap is greater than expected. The resulting p-values were corrected for multiple testing by using the Benjamini-Yekutieli method.

This procedure was also applied to generate Table 1 to study the overlaps of symmetrically and asymmetrically expressed genes in the grafting dataset with previously published sugar-responsive genes.

#### GO enrichment analysis

The gene ontology analysis (GO) enrichment analysis on grafting-specific genes was done with a customized R script using the package GOstats (65). Gene ontology annotation was used from the Bioconductor package org.At.tair.db (67). The p-values calculated by a hypergeometric test were corrected for multiple testing with the Bonferroni correction. A GO category was called enriched if the corrected p-value was below 0.05.

A detailed description, the required data and R scripts to reproduce the clustering (hierarchical clustering and PCA), the statistical analyses regarding the overlap studies, and the GO enrichment analysis are available via the GitHub repository (https://github.com/AlexGa/GraftingScripts).

#### Breaking force measurements

A micro-extensometer was used for all breaking force graft measurements according to a previously published protocol (62). Briefly, grafted Col-0 plants were attached to a moving plate using tough tags (0.94X0.50 inches, white, 679 catalog no. TTSW-1000, DiversifiedBiotech) and a cyanoacrylate glue. The plates were moved apart and the force was measured using a force sensor (Futek LSB200 10g load cell, Futek Inc.). Upon breaking the force dropped. The maximum force measured before breaking was recorded. Images were captured using a Leica SP5 confocal microscope. During the experiment single z-place images were captured and made into a movie in ImageJ.

To measure levels of tissue contamination between grafted top and bottom, grafted green fluorescent protein(*p35S::GFP*) and red fluorescent protein (*pUBQ10::PM-tdTomato*) expressing plants were pulled apart manually and imaged on a Zeiss LSM 700 confocal microscope. Z-stacks comprising the majority of the hypocotyl were made and regions approximately 0.5mm above and 0.5mm below the graft junction were made into projections using the average intensity function on FIJI software, and then the mean intensity quantified for red and green channels on FIJI. The mean intensity of one colour in the tissue tested for contamination was divided by the mean intensity of the same colour in the tissue expressing that transgenes to get a percentage of contamination. A region away from the cut site was also quantified to get a percentage of spectral overlap between red and green channels. The percent spectral overlap was then subtracted from the percentage contamination to get an overall percentage for how much contamination was present.

**Movie S1. Breaking weight measurements at the graft junction, related to figure 1.** A micro-extensometer pulled apart an Arabidopsis plant 9 days after grafting while force was measured using a force sensor and images captured using a Zeiss Sp5 confocal microscope. Here, the transmitted light field is shown. Shoot is on top and root towards the bottom.

**Movie S2. Breaking weight measurements at the graft junction, related to figure 1.** A micro-extensometer pulled apart an *Arabidopsis* plant 9 days after grafting while force was measured using a force sensor and images captured using a Zeiss Sp5 confocal microscope. The same experiment is shown in Movie S1, but here, the GFP and RFP channels are shown. Shoot is on top (*p35S::GFP* expressing) and root towards the bottom (*pUBQ10::PM-tdTomato* expressing).

**Table S1.** Details of previously published datasets used to compare to the grafting datasets.

**Table S2.** Numbers and categories associated with the Bayseq analysis. Categories are defined as grafted (Col:Col), separated (Col_cut) or intact (ungrafted).

**Table S3.** GO analysis for biological process (BP). Shown are the top 20 BP GO terms for the grafting-specific genes. Time point selected are those when there are the most genes in grafted bottom samples (48hrs) or grafted top + bottom samples (120 hrs).

**Table S4.** Normalised reads for all the protein-coding genes in the datasets. By entering the ATG number of interest, a plot is made which shows a differential gene expression profile for the gene of interest.

**Dataset S1.** Details of the Bayesian segmentation analysis providing files containing the ATGs for every cluster for every time point.

**Figure S1.**
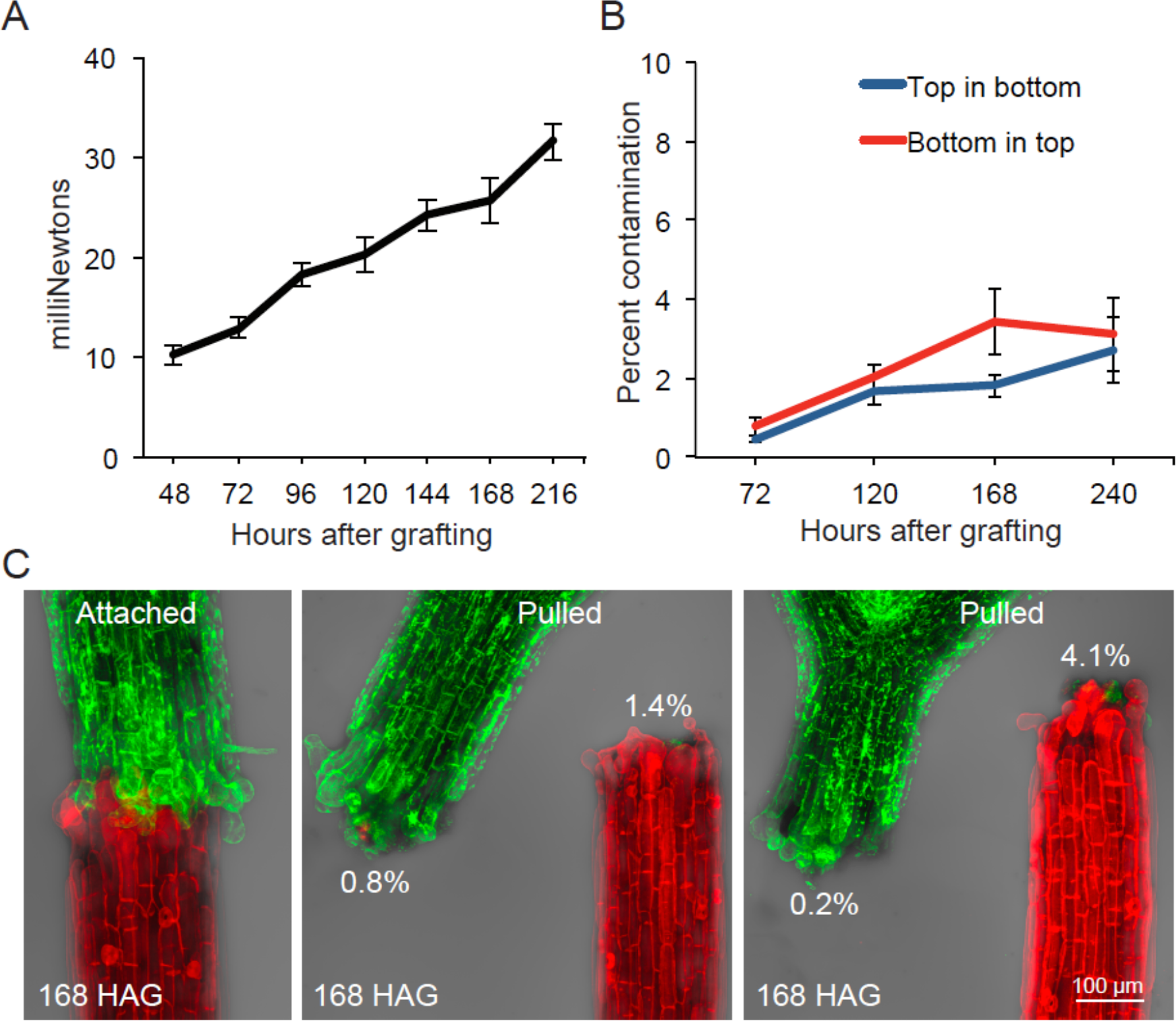
Experimental approach, related to figure 1. (A) The breaking weight required to pull apart grafted plants measured by a micro-extensometer. (B-C) Contamination of top in bottom or bottom in top was less than 4% as measured by grafting green fluorescent protein-expressing plants (*p35S::GFP*) to tomato fluorescent protein-expressing plants (*pUBQ10::PM-tdTomato*) and measuring amounts of fluorescence at the different wavelengths of emission in the top segments relative to the bottom. Images show different plants prior to and after pulling with the percent contamination indicated.

**Figure S2.**
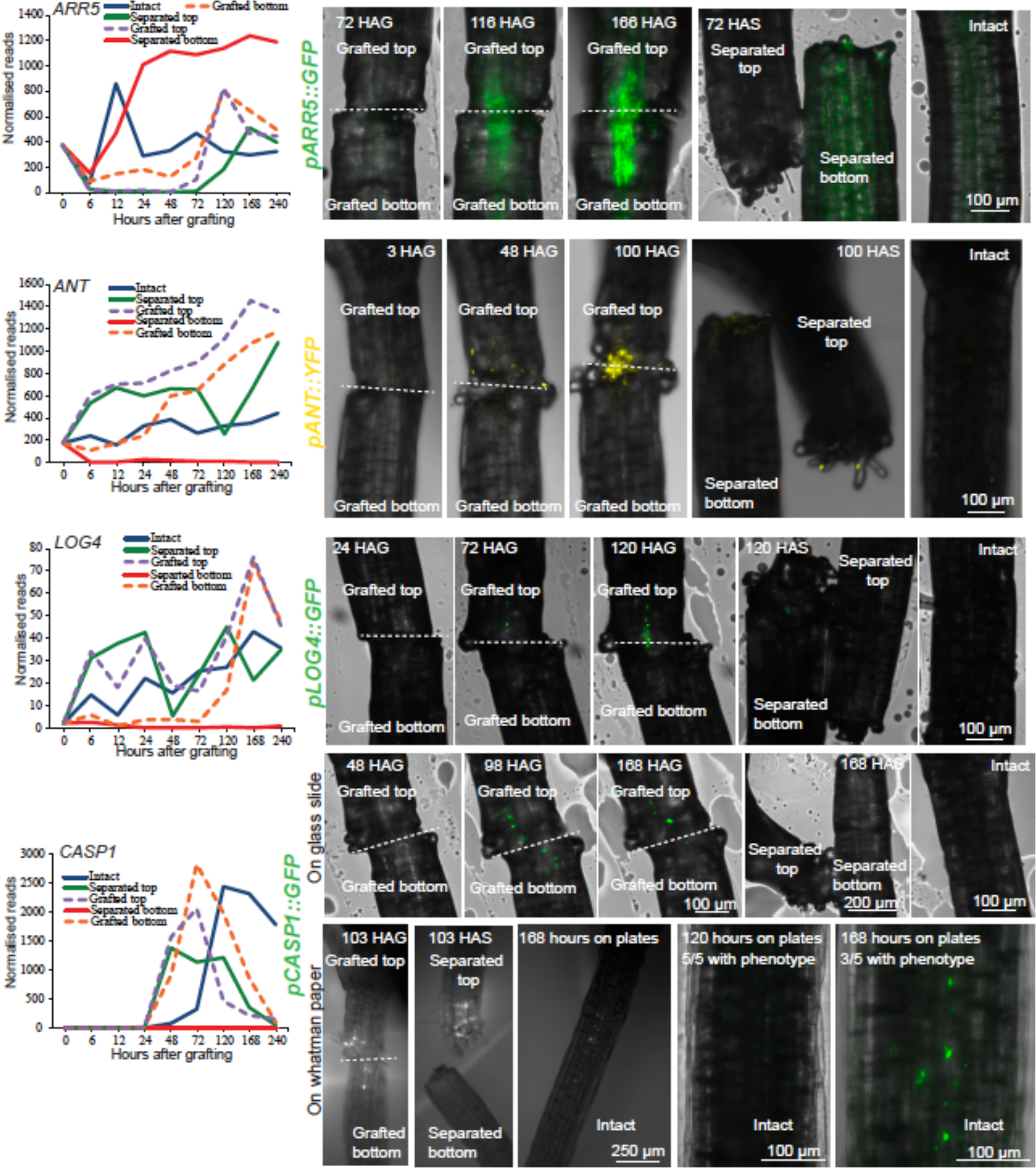
Comparison between RNAseq transcriptome expression profiles and transcriptional fluorescent reporters, related to figure 1. RNAseq expression profiles for various genes upregulated during graft formation were plotted for intact, separated and grafted samples (left panels). Transcriptional reporter-expressing plants were cut and separated, cut and grafted, or left intact. After cutting, plants were imaged and z-projections made at various time points (right panels). For *pCASP1::GFP*, we did not observe a signal in intact plants grafted on glass slides (see Materials and Methods), but observed a signal with 3/5 plants 7 days after grafting on Whatman-nylon membrane, the same condition used for transcriptome library preparation. HAG, hours after grafting. HAS, hours after separation. Dashed lines denote the graft junction.

**Figure S3.**
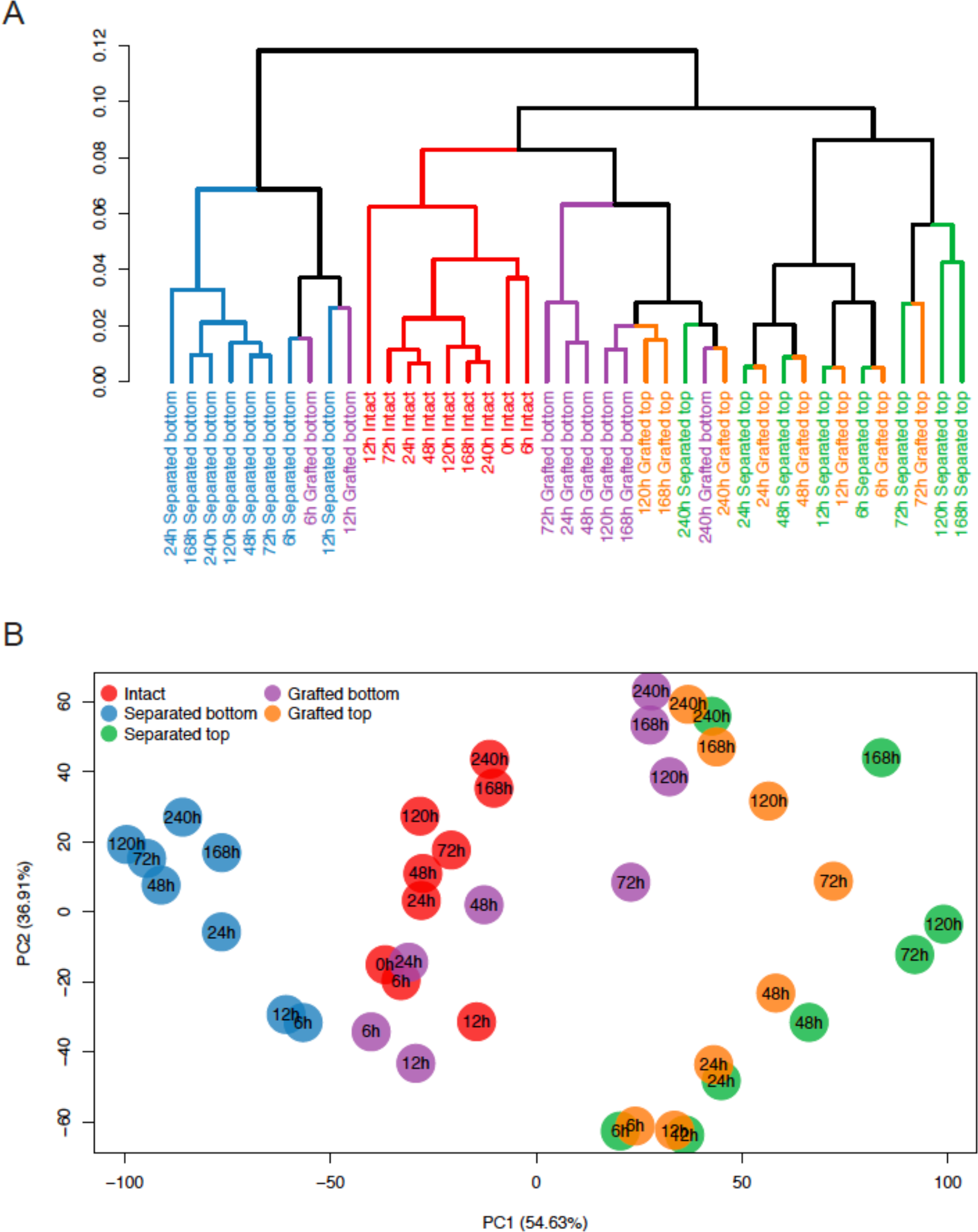
Hierarchical cluster analysis and principle component analysis (PCA) for the transcriptome libraries, related to figure 1. (A) Hierarchical clustering of samples based on log10 transformed TPM values. Similarity between samples was measured by 1 – spearman correlation coefficient. (B) PCA of expression data shows clustering of similar samples. Especially the grafted top and grafted bottom samples are very similar from 120 hours onwards.

**Figure S4.**
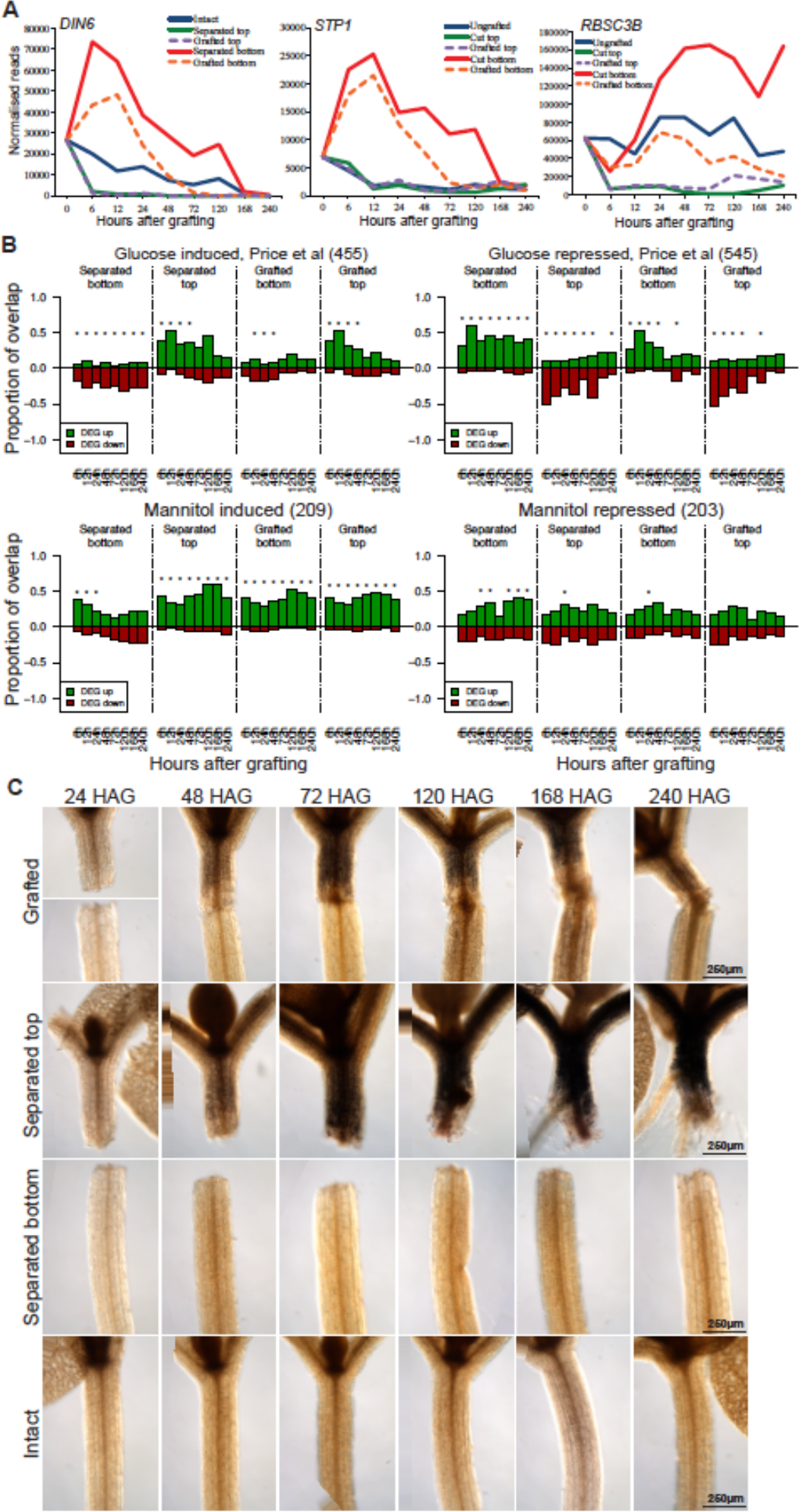
Asymmetry is a feature observed with sugar response and starch accumulation, related to figure 2. (A) Expression profiles for RNAs coded by sugar-repressed genes (*DIN6*, *STP1*) or photosynthetic gene (*RBSC3B*) were plotted for intact, separated and grafted samples. (B) Transcriptional overlap between previously published mannitol-induced, mannitol-repressed, sugar-induced or sugar-repressed genes and our datasets. The numbers in brackets represents the number of glucose or mannitol-responsive genes identified in the previous dataset, and overlap is presented as a ratio out of 1.0 for differentially expressed genes (DEG) up or down regulated in our dataset relative to intact samples. Asterisks represent a significant difference (p < 0.05) between the ratio of up- and down- regulated genes in a previously published transcriptome dataset compared to the ratio of all up- and down- regulated genes in our grafting dataset at a certain time point. (C) Lugol staining of grafted plants at various time points reveals dark brown staining associated with starch accumulation. Upper grafted panels are the same as those present in Figure 2 and shown here to compare with controls. HAG, hours after grafting.

**Figure S5.**
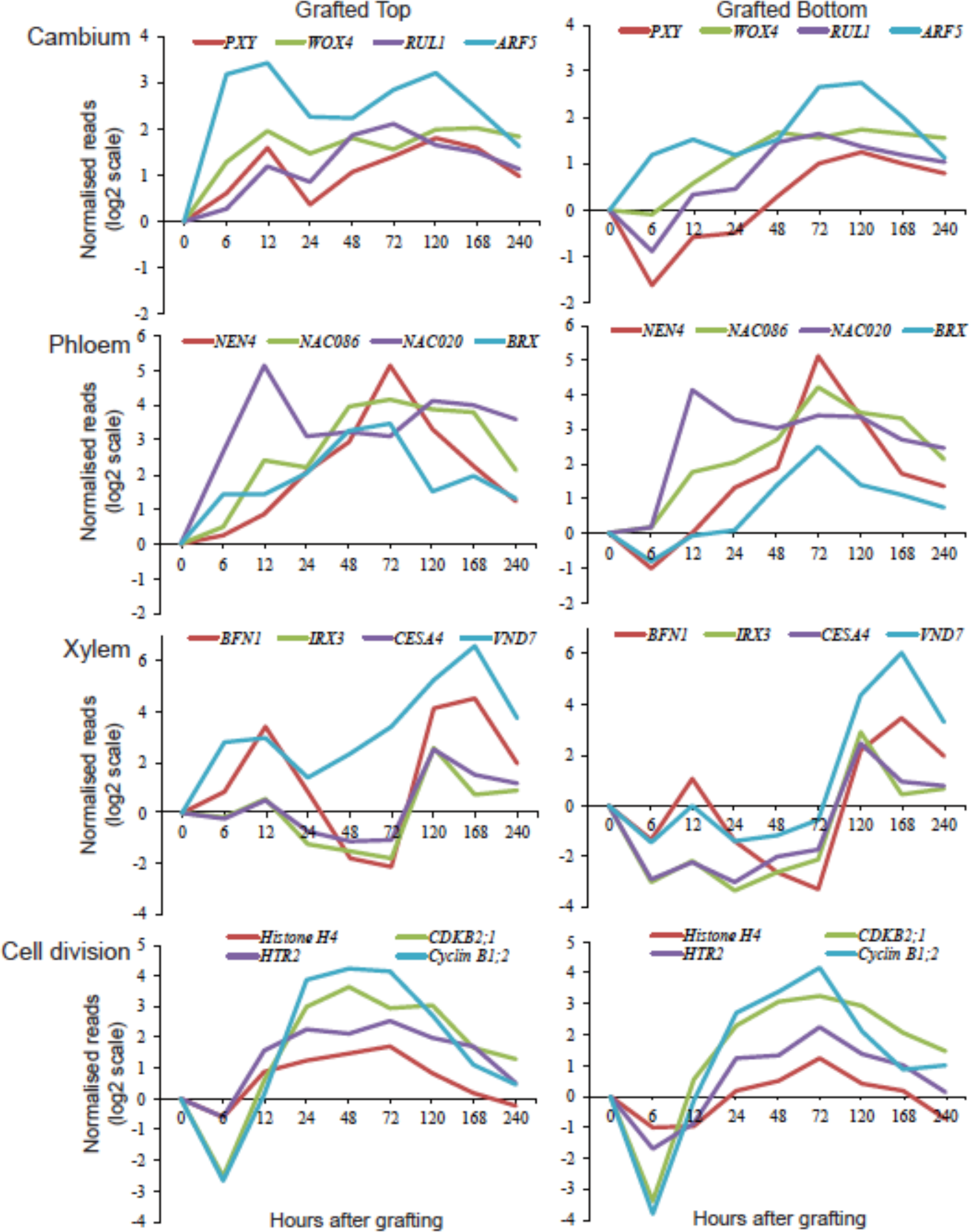
Transcriptional dynamics of genes associated with cambium, phloem, xylem and cell division, related to figure 3. Expression profiles in grafted tops or grafted bottoms for various genes of interest were plotted, normalised to intact samples and plotted on a log_2_ scale.

**Figure S6.**
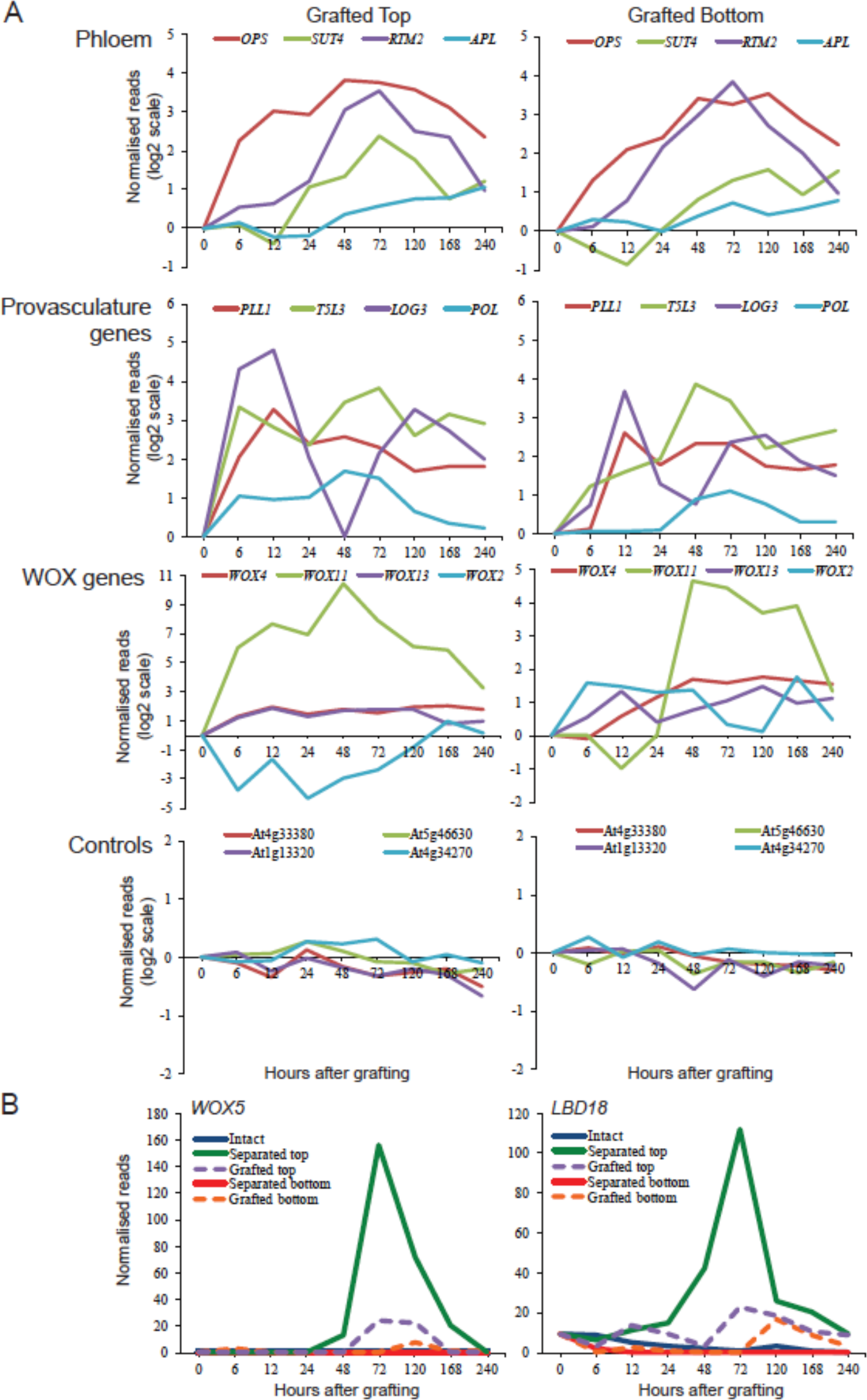
Accumulation of RNA from genes associated with provasculature and phloem, and of RNAs from WOX transcription factors, related to figure 3. (A) RNA accumulation profiles for various genes of interest were plotted over time as measured in grafted tops or grafted bottoms, normalised to intact samples and plotted on a log_2_ scale. The grafting datasets could also be used to investigate the transcriptional dynamics of related genes, such as the sequential activation of WOX transcription factors at the graft junction. (B) Expression levels of a primary root specific transcript (of the *WOX5* gene) or a lateral root specific transcript (from the *LBD18* gene) were plotted over time for intact, separated and grafted samples.

**Figure S7.**
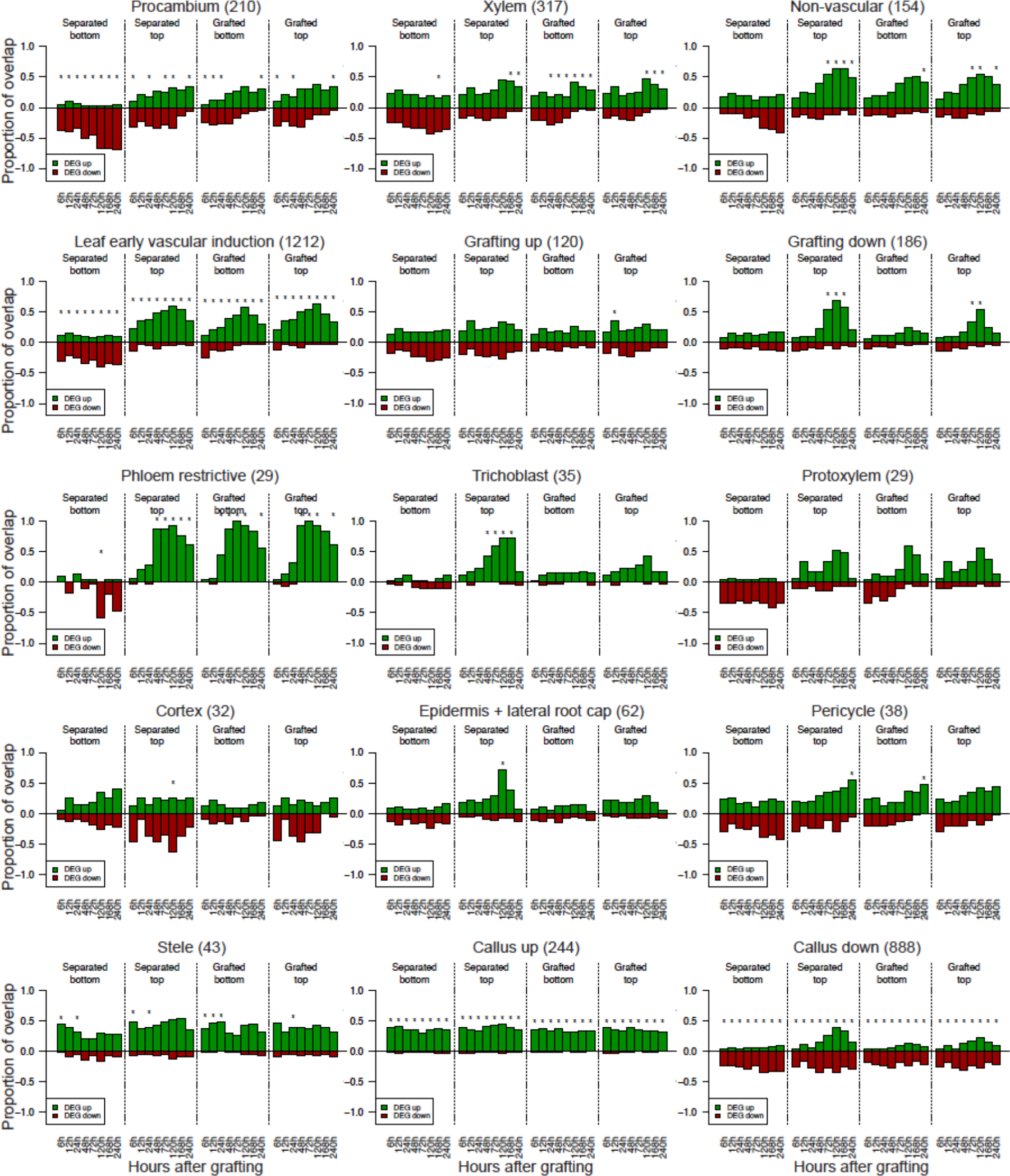
Transcriptional overlap between previously published datasets and the grafting datasets, related to Figure 4. Genes whose transcripts are associated with various cell types or biological processes were taken from previously published datasets (see Table S1) and compared to the datasets generated here. The number in brackets represents the number of cell type-specific or process-specific genes identified in the previous dataset, and overlap is presented as a ratio out of 1.0 for differentiation expressed genes (DEG) up- or down- regulated in our dataset relative to intact samples. Asterisks represent a significant difference (p < 0.05) between the ratio of up- and down- regulated genes in a previously published transcriptome dataset compared to the ratio of all up- and down- regulated genes in our grafting dataset at a certain time point.

**Figure S8.**
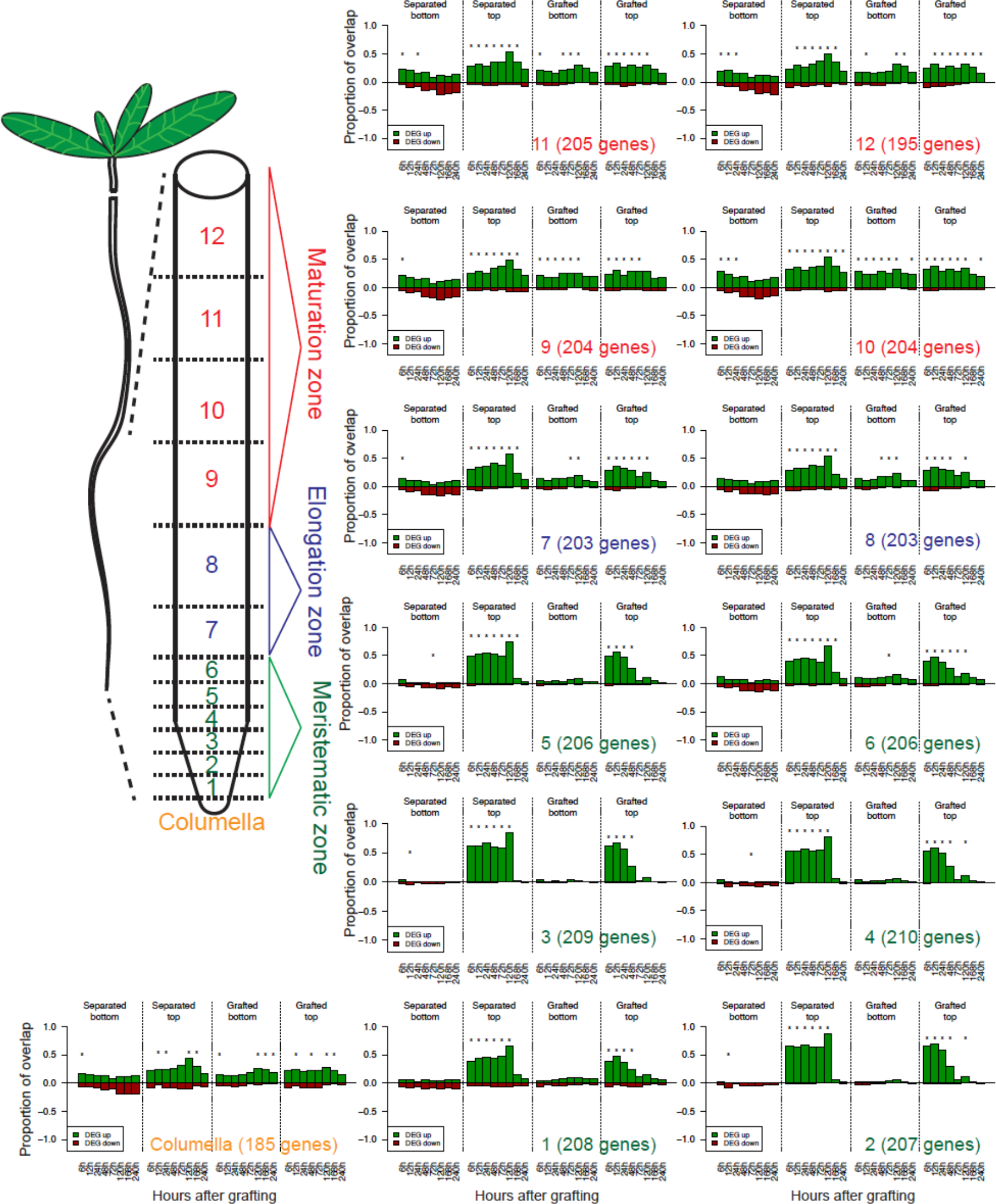
Transcriptional overlap between previously published root transcriptomic datasets and the grafting RNA-seq dataset, related to Figure 4. Genes whose transcripts are associated with various root regions were taken from previously published datasets (see Table S1) and compared to the datasets generated here. Root layer is indicated in the bottom right of each histogram with reference to the cartoon. The number in brackets represents the number of root layer-specific genes identified in the previous dataset, and overlap is presented as a ratio out of 1.0 for differentiation expressed genes (DEG) up- or down- regulated in our dataset relative to intact samples. Asterisks represent a significant difference (p < 0.05) between the ratio of up- and down- regulated genes in a previously published transcriptome dataset compared to the ratio of all up- and down- regulated genes in our grafting dataset at a certain time point.

**Figure S9.**
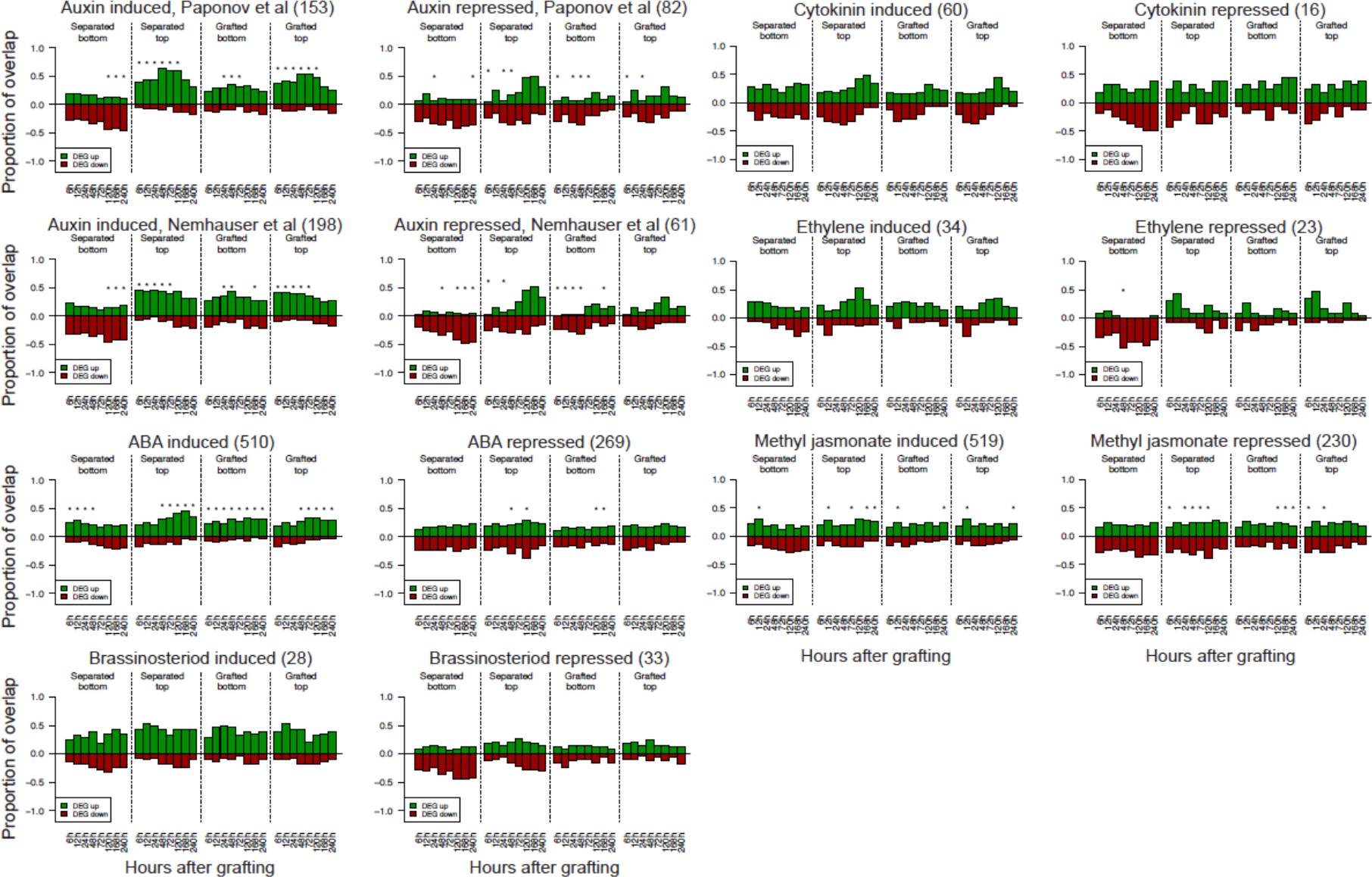
Transcriptional overlap between previously published hormone responsive RNA datasets and the grafting RNA datasets, related to Figure 5. Genes whose differential expression is associated with various hormone responses were taken from previously published datasets (see Table S1) and compared to the genes represented in the RNA-seq datasets generated. The number in brackets represents the number of cell type-specific or process-specific genes identified in the previous dataset, and overlap is presented as a ratio out of 1.0 for differentiation expressed genes (DEG) up- or down-regulated in our dataset relative to intact samples. Asterisks represent a significant difference (p < 0.05) between the ratio of up- and down- regulated genes in a previously published transcriptome dataset compared to the ratio of all up- and down- regulated genes in our grafting dataset at a certain time point.

**Figure S10.**
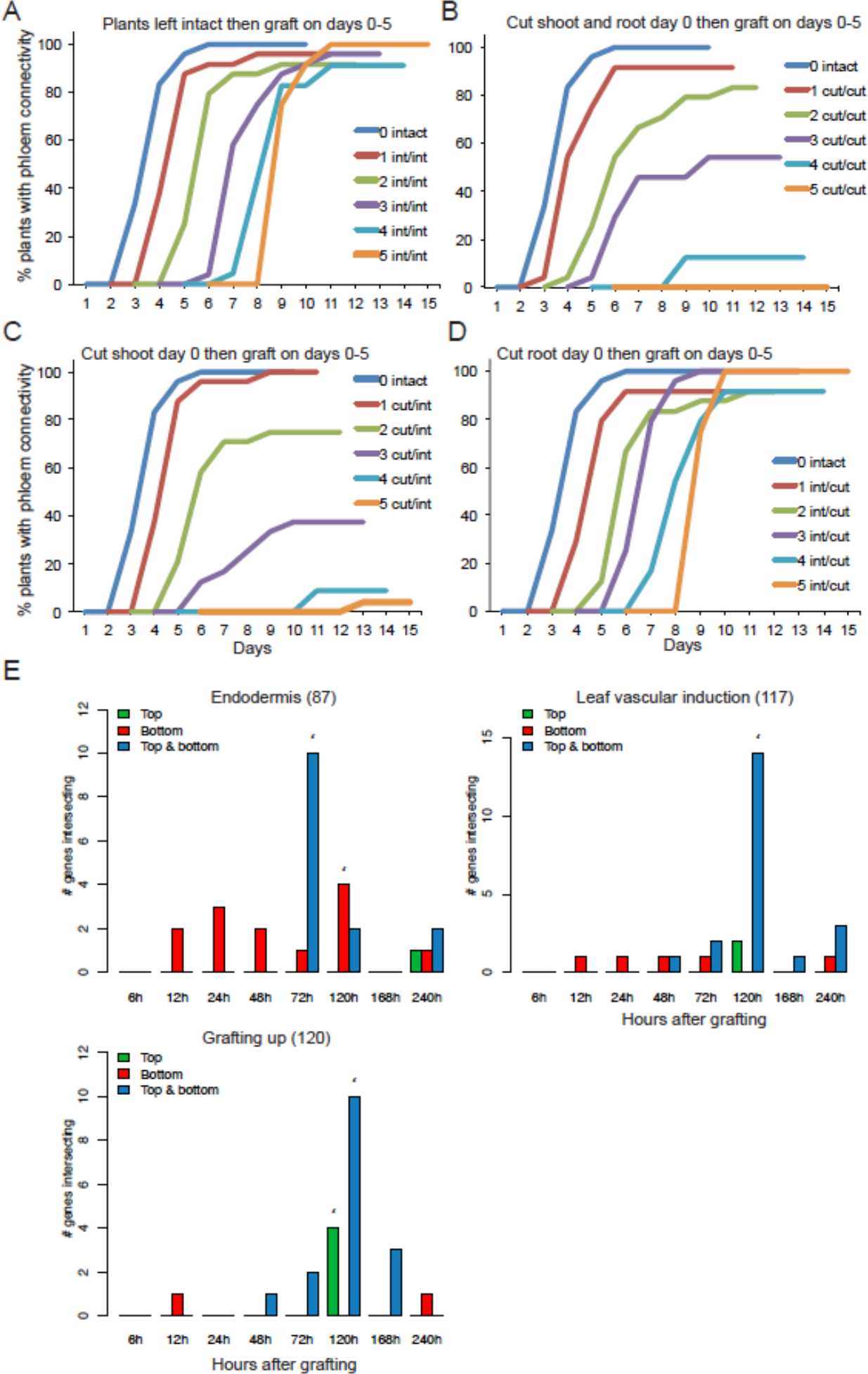
A subset of genes is differentially expressed only during graft formation, related to Figure 7. (A) *pSUC2::GFP* expressing *Arabidopsis* shoots were grafted to Col-0 wild type roots 0-5 days after transferring to grafting plates and kept intact until grafting (intact treatment). GFP movement to the roots for phloem connection assays was monitored over 7 days. (B) Plants were cut at day 0 and grafted at days 0-5 after cutting. (C) Shoots were cut at day 0 and grafted at days 0-5 after cutting. Roots were kept intact until immediately before grafting. (D) Roots were cut at day 0 and grafted at days 0-5 after cutting. Shoots were kept intact until immediately before grafting. (E) Genes differentially expressed in grafted tops and grafted bottoms show overlap with previously published genes whose transcripts are associated with the endodermis, vascular induction and grafting (see Table S1 for treatment information). Asterisks represent significant high overlap (p <0.05) of previously published gene sets that are also differentially expressed in the grafted samples at a certain time point.

